# A modified methyl transferase cofactor to selectively disable gene expression in *E. coli*

**DOI:** 10.1101/2025.04.10.648196

**Authors:** Oliver J. Irving, Samuel Stone, Robert K. Neely, Tim Albrecht

## Abstract

Artificial control of gene expression in bacteria offers interesting prospects for influencing bacterial pathogenicity and antibiotic resistance. We show that the methyl-transferase cofactor, AdoHcy azide, can disable gene expression in modified plasmids in some strains of *E. coli*, where ampicillin and kanamycin resistance as well as eGFP genes were selectively and independently disabled. The disabling of transcription is likely due to steric inhibition during transcription initiation, which is further confirmed by Sanger and nanopore sequencing results. Both sequencing methods showed that 3-6 nucleotides were absent from around the modification site, with the post growth, extracted AmpR/ eGFP plasmid showing evidence of restriction, with sections of the plasmid, including the modification site, missing for the AdoHcy azide modified plasmids. Notably, the AdoHcy azide modification on the DNA is resistant against demethylation in the BL21 strain of *E. coli*.

## Introduction

Epigenetic modification of DNA in bacteria represents a sophisticated layer of regulatory mechanisms that influence gene expression without altering the underlying DNA sequence^1,2^. Unlike in eukaryotes, where epigenetic modifications such as DNA methylation and histone modification are well-characterized, bacterial epigenetics primarily involves DNA methylation and RNA-mediated processes, primarily focused on phage resistance^3^. For instance, the methylation of promoter regions can either repress or activate gene transcription, enabling bacteria to rapidly adapt to environmental changes, regulate virulence factors, and develop antibiotic resistance^4–11^. Understanding and manipulating these epigenetic processes offers exciting possibilities for developing novel antimicrobial strategies, potentially allowing for precise control over bacterial pathogenicity and the mitigation of resistance mechanisms^12^.

Recent advances in synthetic biology and genetic engineering have provided researchers with powerful tools to manipulate bacterial genomes with unprecedented precision^13^. Techniques such as CRISPR-Cas, synthetic promoters, and inducible expression systems allow for the fine-tuning of gene expression^13–16^. These technologies offer a new paradigm in the fight against bacterial infections. However, they are highly specialized, requiring dedicated synthetic biology laboratories to carry out. RNA interference (RNAi) and short-hairpin RNAs (shRNA) techniques offer alternatives to disable genes, targeting specific mRNA sequences and preventing their translation^17–21^. However, they can target non-specific sequences and affect the expression of other genes^22^. Once the affecting RNA is introduced, the target gene is knocked down until the RNA pool has been depleted or complementary oligonucleotides to the RNA strands are transfected and gene expression restored^23,24^.

In our previous work, we showed how methyl transferase-based AdoHcy azide (AHA) modification of dsDNA can prevent T5’ exonuclease mediated digestion of short dsDNA fragments and inhibit PCR amplification^25^. We hypothesized that this effect was likely due to steric inhibition of exonuclease and polymerase activity on the DNA strand, respectively. This raised the question whether the same approach could be exploited to control DNA enzyme interaction *in vivo*, potentially as a new “chemical” way of controlling gene expression. This would substantially widen the toolkit available for this purpose, not only due to the wide variety of structural variations accessible through synthetic chemistry, but also new functional paradigms, such chemical or photochemical switching of gene transcription.

In the present paper, we demonstrate the successful implementation of such an approach. As shown in Fig. 1, we tested plasmids with different modification inserts (no insert, unmodified insert, and CH3-/MTC5-/AHA-modified inserts, plasmids 1-5) and different functional “reporter” genes (AmpR, KanR, and AmpR + eGFP, in plasmid 6), which were transfected into two strains of *E. coli*, D5Hα and T7-enhanced BL21. The latter strain retains an intact base excision repair (BER) pathway, which may recognize and remove certain modified bases if they are perceived as DNA damage. Non-transfected cells and heat-treated cells (without transfection) were employed as further controls. Hence, this study design enabled testing the impact of increasing size of the DNA modification on enzyme function; to demonstrate the effect of the AHA modification in targeting individual genes; and show the specificity of gene expression control (via the 2 gene encoding plasmid 6).

**Figure 1.**
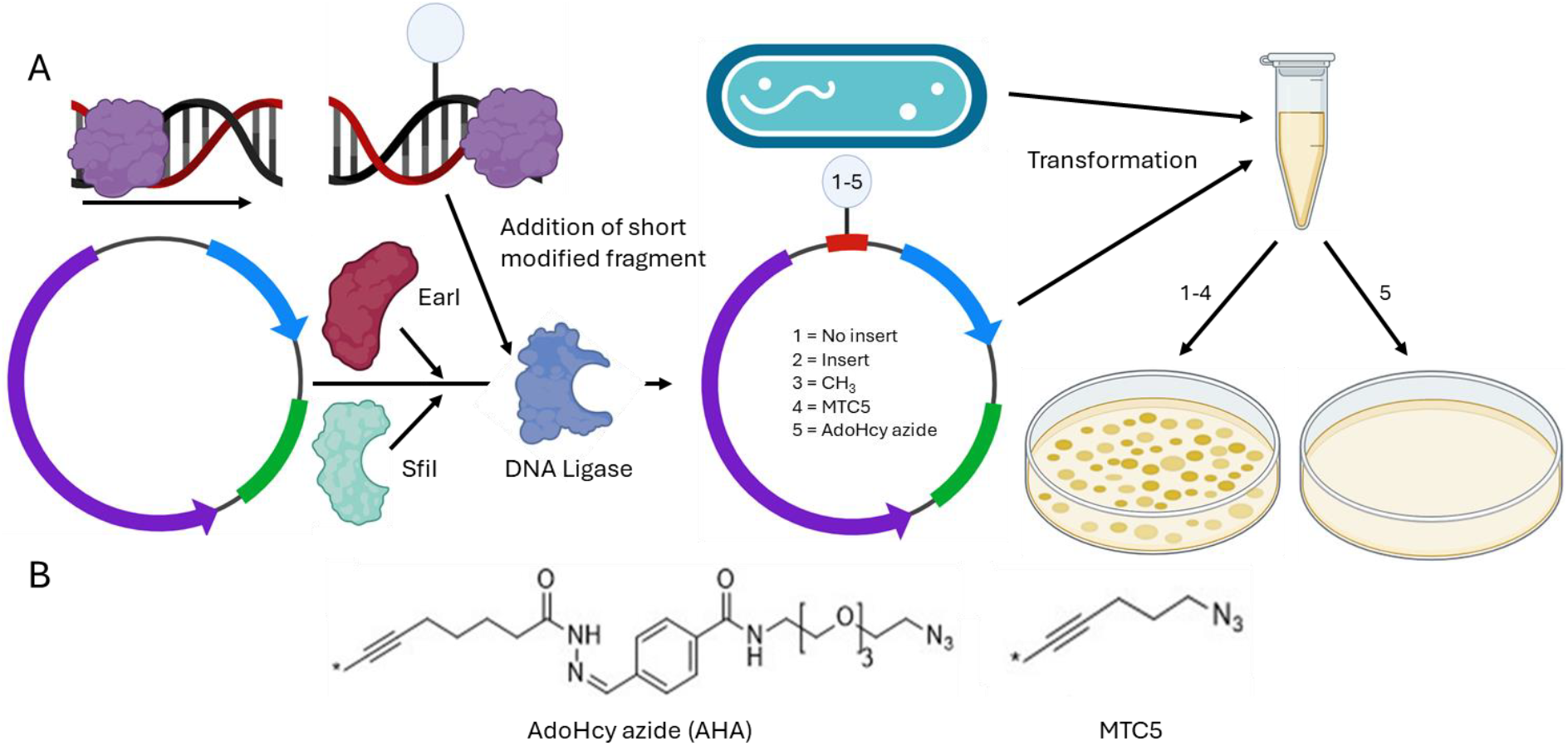
Plasmid design and modification. A) Modification groups are transferred to the short dsDNA ligation insert strands (red) using M.TaqI. S10-pMA-RQ (AmpR) is restricted using R.EarI and R.SfiI to remove additional DNA sequence (blue segment: resistance gene, green: origin, purple: S10 custom gene). The modified ligation insert is then added to the restricted plasmid with a DNA ligase to incorporate this into the plasmid backbone. The assembled plasmid and *E. coli* are subsequently combined in SOC media for heat shock transformation. The bacteria are then plated and allowed to grow on ampicillin-or kanamycin-containing media, where bacteria with plasmids 1-4 continue to express the resistance gene and grow, while plasmid 5 does not. B) The chemical structures of the synthetic modification groups AHA and MTC5.

## Results

As noted above, two plasmid designs contained a single antibiotic resistance gene, AmpR or KanR, and a third with both AmpR and eGFP genes separated by a non-coding, spacer region of DNA (plasmid 6). Where applicable, spacer sequences (red) were added at the target promoter site, without disrupting the native promoter sequence, as confirmed by nanopore sequencing for plasmids 1 and 2, see below and supplementary information (SI) section 2.

Some inserts were modified using M.TaqI and different cofactors (blue circle, 3-5) before incorporation into the restricted plasmids, see Methods. Note that M.TaqI modifies dsDNA predominantly on one of the two strands, which is relevant to our discussion below^26–29^. The structures of the transferable linkers from the AHA and the MTC5 cofactors are depicted in panel B^26^. The bacteria were transformed using the modified plasmids grown. Accordingly, modification inserts that disable gene expression would lead to a lack of antibiotic resistance or fluorescence in the bacteria, despite the presence of the relevant genes, while those without inhibitory effect would still display gene-specific responses. Statistical testing was performed using two-factor ANOVA (α = 0.05), see SI, section 5, for further statistical analysis.

This expectation is indeed borne out in colony growth experiments, as summarized in Fig.2A and in the SI, section 1. Table 1 shows the results obtained for the two bacterial strains, DH5α and BL21 (left), three different plate conditions (no antibiotic/control, ampicillin, or kanamycin present) and the plasmid type. Colony growth is indicated by a tick, lack thereof by a cross/red shading, as determined by statistical analysis (two-way ANOVA, α = 0.05, SI4).

**Figure 2.**
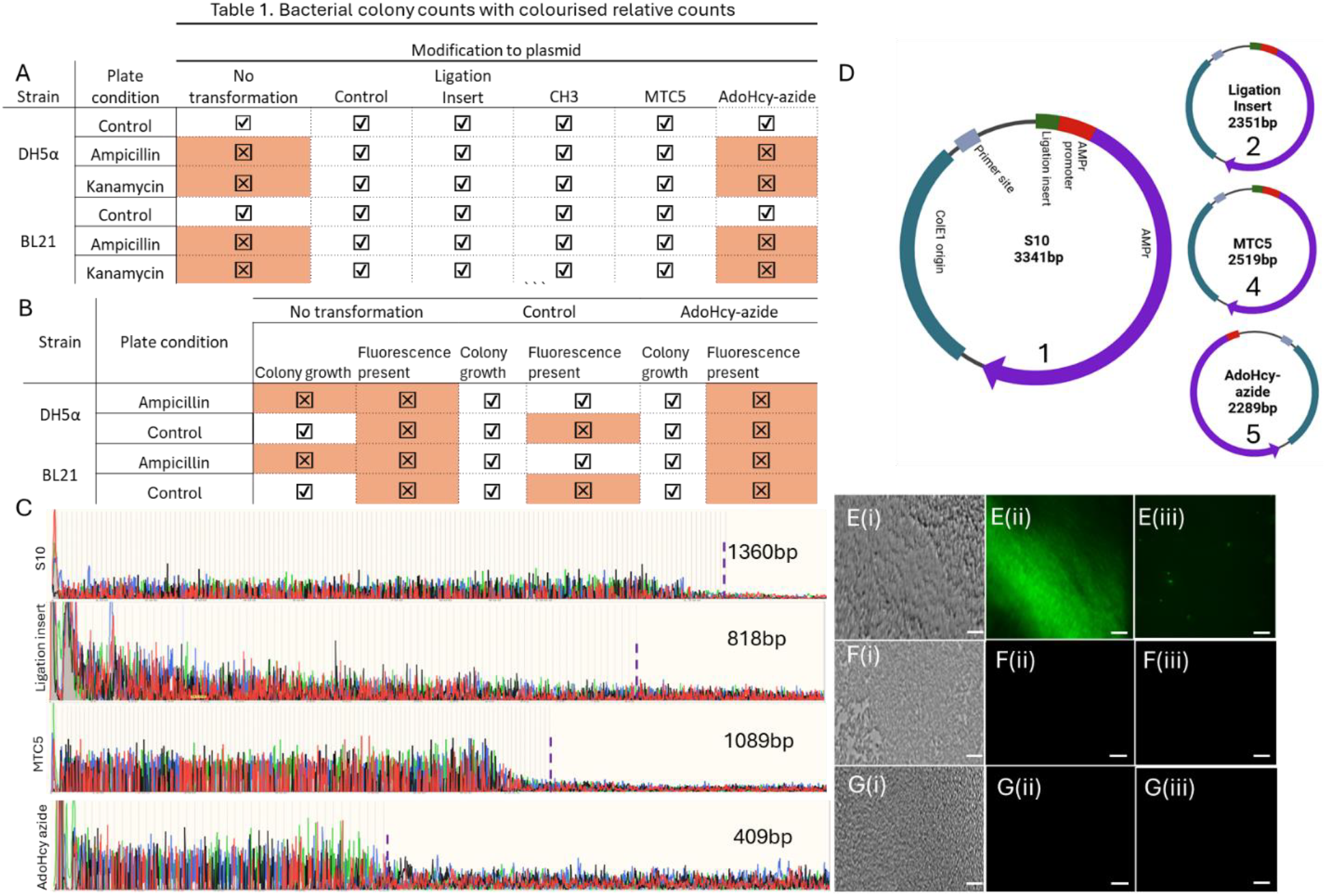
A) Table showing presence or absence of colony formation on control, ampicillin and kanamycin containing plates. Results for no colony formation highlighted in red. (B) A table showing the presence or absence of colonies and fluorescence on control and ampicillin containing plates. (C) Chromatogram results from Sanger sequences showing plasmids 1,2,4, and 5, where maximum read lengths are highlighted (purple dashed line). (D) Illustration of the ON sequencing results. The green highlight region on plasmid 5 is absent, indicating that it could not be sequenced. Control BL21 containing the unmodified EGFP plasmid (E) were imaged using brightfield (i) and green fluorescence (ii) channels, and single cell fluorescent images (iii). (F and G) DH5α and BL21, respectively, containing the AHA modified plasmid 6 showing strong bacterial numbers (i), however in the green fluorescent channel only background could be measured with no discernible features (ii and iii). Scale bars represent 10µm.

Notably, bacterial growth is detected for all plasmids 1-5 in the absence of ampicillin or kanamycin and while there was a small decrease in colony count when comparing transformed and non-transformed bacteria (see SI4), we found no statistically significant difference in colony count between the different plasmids (two-way ANOVA, p = 0.75, 0.99, 0.99, 0.12, with respect to plasmid 1, SI4). This suggests that the presence of the modified plasmids does not adversely affect initial bacterial seeding and growth in control conditions. In the presence of the respective antibiotic on the growth plates, no colonies could be found for bacteria without the plasmid, since those lack the relevant resistance gene and perish. On the other hand, bacteria with the plasmids 1-4 did grow under the same conditions and in both bacterial strains, suggesting that the relevant resistance gene was not only present but functional. Finally, bacteria transformed with the AHA-modified plasmid 5 did not grow in the presence of the antibiotics, even in extended growth assays for up to seven days, in either DH5α or BL21. In principle, this could either mean that the relevant plasmid, and hence the resistance gene, are not or no longer present in the bacteria or that the gene is present but cannot be transcribed, resulting in a lack of antibiotic resistance and bacterial death. Experiments with bifunctional plasmids addressed this question and showed that the latter is indeed the case. To this end, bacteria were transformed with plasmid 6, containing both AmpR and eGFP genes. In unmodified plasmid 6, we expect both eGFP and AmpR transcription to be active, resulting in green fluorescence emission from GFP, and survival of the bacteria on the ampicillin containing plate. After modifying the eGFP promoter with AHA, the eGFP gene would be disabled, and no GFP fluorescence detected. However, with AmpR transcription still active the growth of the bacteria would be unaffected on ampicillin containing plates.

Results for colony growth and fluorescence emission show that this is indeed the case, Fig.2B and 2E-G. Control plates with untransformed bacteria did not grow. BL21 transformed with the unmodified control plasmid showed no statistically significant difference in the number of colonies, compared to the ampicillin-free control plate (p = 0.37), but strong fluorescence emission from GFP, Fig.2E. Correspondingly, for BL21 transformed with the AHA-modified plasmid, there was again no statistically significant difference in colony growth compared to the control condition (p = 0.17), while GFP fluorescence is now absent, Fig.2F. This was also observed in the DH5α, Fig.2G. In line with our hypothesis, this not only confirms that the transformed plasmid is present and active, in terms of AmpR transcription, but also that the eGFP gene can be disabled in the bacterial population due to the presence of the AHA modification in the initial plasmid. Moreover, the latter appears to be robust against passive enzymatic demethylation in BL21, under the experimental conditions used.

In order to gain sequence-level information and hence further insight into the mechanism of enzyme inhibition, we applied both Sanger and nanopore sequencing to plasmids 1, 2, 4, and 5. It is worth noting here that both sequencing methods rely on enzymatic processing. Sanger sequencing involves a DNA polymerase, and we had previously found that AHA modification can suppress the action of DNA polymerase in PCR amplification^25^. Nanopore sequencing on the other hand, relies on helicase-based unwinding of the DNA strand prior to readout in a narrow pore channel. It is therefore interesting to compare the results from both methods, as this will also provide further insight into DNA enzyme interaction with AHA. Taking the Sanger sequencing results first, the (mono-directional) primer end was positioned 412 bp from the modification site, and the results are shown in Fig.2C (full sequence information in SI2). The unmodified AmpR plasmid 1 could be read to a length of 1360 bp (maximum read length) with a quality (Q) of 42, plasmid 2 could be read to 818 bp (Q = 15), and plasmid 4 to 1089 (Q = 48). In all cases, the read sequence included the modification site and accurate sequence information could be obtained. However, plasmid 5 could only be read to 409 bp (Q = 42). It was not possible, over several sequencing runs (n = 3), to sequence past this point as indicated by the sharp drop off in the chromatogram, Fig.2C (bottom). This indicates that the presence of the AHA group likely prevented the polymerase enzyme from passing over the modification site and is in-line with our previous findings, namely that AHA modification inhibited PCR based DNA amplification^22^.

In the case of nanopore-based sequencing, AmpR plasmids 1, 2, as well as the MTC5-modified plasmid 4 were all sequenced as expected when compared against the manufacturer provided plasmid 1 sequence. However, sequencing of plasmid 5 unexpectedly produced an inverse read instead, as illustrated in Fig.2D (full sequence information in SI2)^30^. Observing the inverse read is consistent with only one strand dominating the read to generate the final sequence ^31^.

This indicates that the modification cannot pass through the pore, and only the unmodified strands can contribute to the read. To this end, we found 6-bases missing ahead of the modification, compared to the Sanger results 3 missing bases, indicating that inhibition occurs further from the reading site. Finally, the methyl transferase site is absent from the sequence, and the AmpR promoter has also been shortened by 3 bases, cf. SI3. This raised the question as to how the expression of the relevant genes is prohibited given that the AHA modification cannot be preserved during cell division.

Nanopore sequencing of the AHA modified plasmid 6 extracted from bacteria post-growth, as shown in SI2, revealed that the plasmid is in fact shortened by 1944 bases. The eGFP gene is missing a proportion of its promotor region, including the modification site, and a large section of spacer DNA between the two genes has also been removed. This explains why the eGFP gene is no longer expressed, while antibiotic resistance is unaffected. With a reduced yield from purification (25ng/μL for the modified, and 251ng/μL for the unmodified plasmids), we suggest that during replication, the inability to copy the modification site initiates a recombination event, removing parts of the plasmid and annealing the remaining sections of the plasmid to keep the AmpR gene intact (modification-induced DNA restriction and repair event)^32,33^. This is supported by sequencing results in that the remaining sequence matches the unmodified plasmid.

In conclusion, we have shown that chemical modification of DNA at the relevant promoter site via the synthetic methyl transferase cofactor AHA can be employed to disable gene expression in *E*.*coli* effectively and selectively, using AmpR, KanR and eGFP as reporter genes in two bacterial strains, DH5α and BL21. The AHA modification appears to resist passive demethylation in the BL21 strain of *E*.*coli*. We suggest that recognition of the AHA initiates restriction of the eGFP plasmid, reducing its size by 1944 bases, but leaving the AmpR gene intact. The inhibition effect was not observed for the CH3 and MTC5 modifications, suggesting steric inhibition of enzyme action may play role in the case of AHA. Most importantly, however, our findings open the door for a new methodology to control gene expression in vivo, in *E*.*coli* and in other target systems, by fine-tuning the chemical structure of the cofactor, potentially making it selective to specific DNA or RNA processing enzymes, or by introducing switching capabilities, such as photo-induced isomerisation or responsiveness to environmental stimuli inside the cell.

## Methods

All bacterial work was carried out under aseptic conditions on a 70% ethanol sterilized surface, next to a Bunsen burner, with all equipment sterilized using ethanol solution and flame. All enzymes and reagents were obtained from NEB (Ipswitch, USA) unless otherwise stated. Post modification or assembly step, proteinase K and heat deactivation (70°C) was used to prevent enzymes from entering the next assembly step. In short, the method is described here and illustrated in figure 1 (detailed methodology in SI1). The custom plasmid S10-pMA-RQ (AmpR), containing 1000bp of filler DNA between 2 SfiI sites, was used for all ampicillin resistance experiments. The plasmids used for kanamycin resistance and eGFP were pET-28c(+) and pCSDEST2-EGFP respectively.

A short spacer sequence (I), containing the AmpR promoter and a TCGA site included prior to the promoter, was modified to contain a CH3, MTC5, or AdoHcy azide group using M.TaqI methyl transferase. The plasmid was restricted using the EarI and SfiI sites. Four separate plasmids were prepared to contain each modification, a control, and an unmodified sequence inserted plasmid. Spacer sequences were added at 3X concentration and ligated using a quick ligation kit (NEB). This was repeated for the KanR and eGFP plasmids as well. Reassembled plasmids of interest were sequenced using Sanger and Oxford Nanopore sequencing (SI2). *E*.*coli* strains DH5α and T7 enhanced BL21 were transformed using the heat shock method to contain the modified, re-ligated plasmids and plated on ampicillin containing LB agar plates. The plates were incubated at 37°C for 48 hours and the number of colonies counted using Fiji-ImageJ. Unmodified and modified plasmid 6 were extracted from cells post growth using QIAprep® Miniprep kit (QIAGEN).

To ensure that AdoHcy-azide did not affect bacterial growth, antibiotic free plates were seeded with untransformed *E*.*coli* and incubated with saturated filter disks containing 100X the working concentration of each modification group. A zone of inhibition was not observed at any of the incubation sites. This indicated that the reactant themselves did not induce bacterial death, and that any growth suppression observed would be due to the DNA modifications.

## Acknowledgements

The authors would like to thank Professor Willem van Schaik, University of Birmingham, for reading the manuscript.

## Author contributions

O.J.I and T.A conceived the study; O.J.I developed the protocol and performed experiments with help from S.S; O.J.I performed analysis. The manuscript was written through contributions from all authors. All authors have given approval to the final version of the manuscript.

## Additional Information

The following files are available free of charge.

Supporting Information with further details on plasmid preparation protocols, bacterial growth assays, gel electrophoresis, plasmid sizing, Sanger and Nanopore sequencing information, and statistical analysis, not included in the main manuscript.

