## Supporting Information for "A modified methyl transferase cofactor to selectively disable gene expression in *E. coli*"

#### SI1. Methodology

All bacterial work was carried out under aseptic conditions on a 70% ethanol sterilized surface, next to a Bunsen burner, with all equipment sterilized using the ethanol solution and flame.

**Plasmid modification**

Plasmids were modified using restriction, and ligation insertion of a short dsDNA fragment. The insertion sequence is provided below. The modification method for the short dsDNA fragments is reported in Irving *et al*[22].

Short dsDNA plasmid inserts (500 ng/µL, sequence below, was incubated with 2 µL Cutsmart buffer 10X, 1 µL of the modification donor at 100 µM ($CH_3$ (SAM), MTC5 (SAH-MTC5), or AdoHcy azide (SAH-AdoHcy azide, AHA), illustrated in the main paper, and used independently), 0.5 µL m.TaqI (NEB) and 15.5 µL of nuclease free water (Merck), adding the enzyme last. This was incubated at 40°C for 1.5 hours. To this, 0.5 µL of proteinase K (800 units/mL, NEB), was added and incubated at 40°C for 1 hour. This was allowed to cool to room temperature for 20 minutes post heating. The solution was then purified using a PCR clean-up kit (GenElute™, Sigma-Aldrich), and stored at 4°C until required.

5’ - TAG TCG AAT CGA GTA CCG - 3’

5’ - TTT CGG TAC TCG ATT CGA – 3’

To insert this sequence into the plasmid (S10-pMA-RQ, pET-28c(+), and pCSDEST2-EGFP) at the gene promoter site, a 50 µL digestion was set up using the parameters described in table S1, and incubated at 37°C for 1 hour, before incubation on ice.

**Table S1. EarI restriction reagents.**

| Reagent | Concentration |
|---|---|
| DNA | 1 µg |
| 10X rCutSmart Buffer | 5 µL (1X) |
| EarI | 1.0 µL (20 units) |
| Nuclease-free Water | to 50 µL |

To reassemble the plasmids to contain the modified insert, in a new vial, 50 ng of cut plasmid, 150 ng of modified insert, and 5 µL of Quick Ligation™ Kit (NEB) were added and incubated at room temperature for 15 minutes. Plasmids were stored at -20°C until transformation.

**Transformation**

Competent *E.coli* cells (DH5α and T7 enhanced BL21, Thermofisher Scientific and NEB respectively) were removed from storage at -80°C and allowed to thaw on ice for 30 minutes. 50 µL of the thawed cells were placed into a new vial and 3 µL of each respective plasmid DNA was added and incubated on ice for 30 minutes. Each vial was then heat shocked at 42°C water bath for 60 seconds. The vials were placed back on ice for 2 minutes before 1 mL of SOC media was added to each vial and incubated on a 37°C shaking incubator for 45 minutes.

**Bacterial growth plates**

Three bacterial plate conditions were used to determine the presence, and effect, of the plasmid modification on bacterial growth. These conditions were the inclusion and absence of ampicillin and/or kanamycin as a selection factor. Where appropriate these are noted in the results section. LB agar was created using 40 g/L of LB agar broth (Thermofisher) and distilled water, and autoclaved for 30 minutes. Where appropriate, 100 mg/mL Ampicillin sodium salt (Thermofisher) or 50 mg/mL kanamycin (Merck) was added to the LB agar, once the temperature had reached 50°C. 15 mL of the prepared agar was poured into 7.5 cm petri dishes. The plates were allowed to cool to room temperature for 45 minutes until the agar had completely solidified. 30 µL of each of the transformed bacteria was pipetted onto the agar plate and swept across the surface using a sterilized glass spreader. The plates were incubated at 37°C for up to 48 hours. Growth was determined by the number of colonies.

**Bacterial colony growth, examples**

The growth plates examples, as shown in figure S1, illustrate the colony formation observed after transformation. The number of colonies were counted using the circular finding tool in Fiji-imageJ and checked manually to ensure reliability. The labelling on the plates are: "Con" for no plasmid, "S10" (plasmid 1, as defined by the main paper), "I" (plasmid 2), "$CH_3$" (plasmid 3), "MTC5" (plasmid 4), and "AW" (plasmid 5).

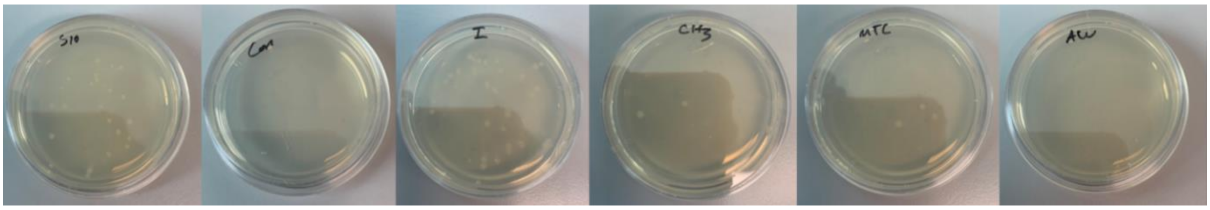


Figure S2. Example ampicillin containing LB agar growth plates for each transformation condition.

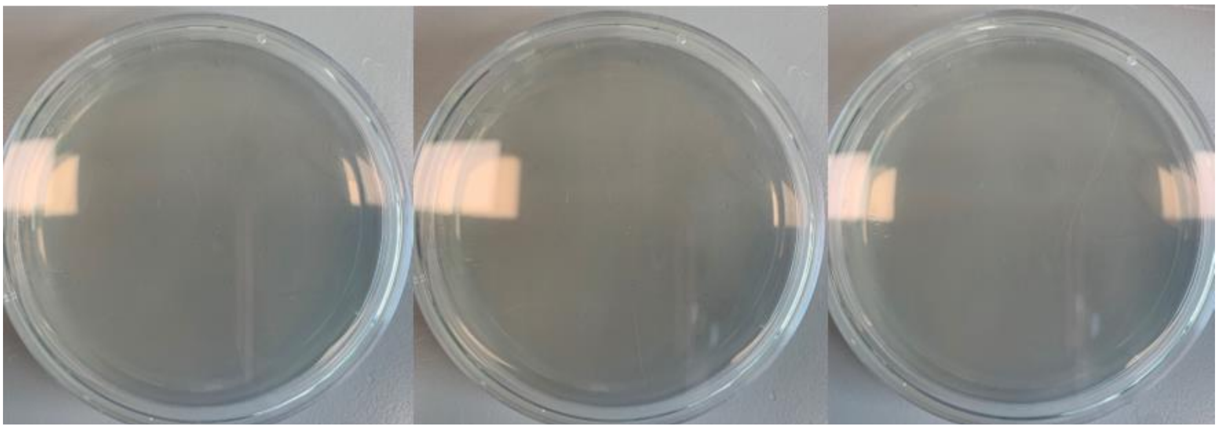

Figure S1. Ampicillin contain plates for 3 separately prepared bacterial samples transformed using AdoHcy azide, imaged after 1, 3, and 7 days (left to right) of growth at 37°C.

The growth assays were performed 3 times. Transformed bacteria containing plasmid 5 were moreover subjected to an extended growth assay over 7 days, figure S2. The results of this assay show that there is no colony formation, indicating that the bacteria cannot grow due to the presence of the ampicillin.

**Imaging**

Bacteria were sampled from plate colonies and swabbed on a standard glass microscope slide, with a cover slip added. Plate images were taken using a handheld camera. For single cell images, bacterial swabs were diluted in 1 mL of SOC media, with 20 µL pipetted onto a microscope slide and a cover slip added. All slides were imaged using a Nikon Ti2 Eclipse under brightfield and green fluorescence illumination. Internal calibration (1 px = 0.108 µm) was used for all measurements and size calibrations.

**Plasmid sizing**

Modified and reassembled plasmids were purified using a PCR clean-up kit (GenEluteTM, Sigma-Aldrich). The products were then run on a 1% agarose gel at 75 V for 40 minutes. To 100 mL of 1X Tris-acetate-EDTA buffer (TAE, Tris (hydroxymethyl) aminomethane 1 g, EDTA, tetrasodium < 1 g, Water 98 mL, Disodium EDTA ~1 g, 1,3-Propanediol, 2-amino-2-(hydroxymethyl)-, acetate (salt) ~1 g, Fisher Scientific), 1 g of agarose powder (Sigma-Aldrich) was added. This was microwaved for 2 minutes until the agarose had completely dissolved. The solution was allowed to cool to 50°C then poured into a gel tray with a 15 well comb in place and incubated at room temperature for 30 minutes for the gel to set. To 5 µL of each DNA sample, and DNA ladder (GeneRuler 1 kbp plus DNA Ladder, ThermoFisher), 1 µL of 6X DNA loading buffer (ThermoFisher) was added and homogenised. The gel was then placed into a Mini-Sub Cell GT Cell (BIORAD), and this filled with 1X TAE buffer until the gel was covered. The well comb was then removed and the first well filled with 5 µL of a DNA ladder (GeneRuler 1 kbp DNA Ladder, ThermoFisher). The remaining wells were filled with 5 µL of the DNA samples under analysis or DNA standards. The gel was then placed in 1X Sybr Gold (ThermoFisher) for 45 minutes before imaging in a UV illumination box (BIORAD). The gel images were then analysed using Fiji-ImageJ.

Gel results show that the expected size of the initial plasmids matches the manufacturer specifications and the modifications do not appear to affect the integrity of the plasmids for transformation.

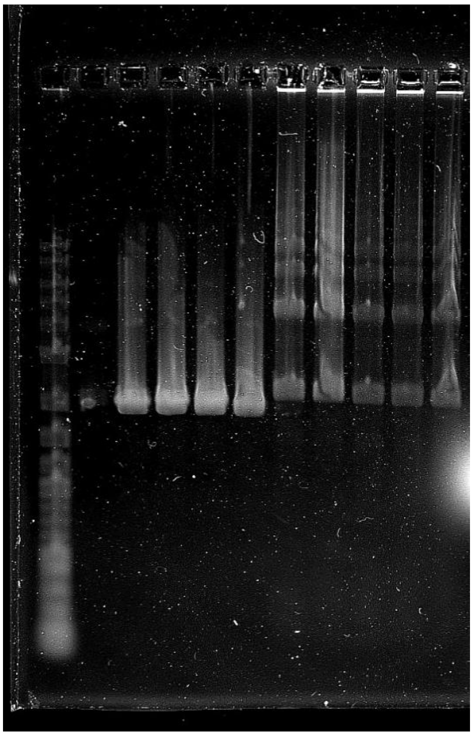

Figure S3. Gel image showing the assembly of modified plasmids (lane 1: gene ruler (1kbp plus, Thermofisher), lane 2: AmpR plasmid 1, lane 3: AmpR plasmid 2, lane 4: AmpR plasmid 3, lane 5: AmpR plasmid 4, lane 6: AmpR plasmid 5, lane 7: eGFP plasmid 1, lane 8: eGFP plasmid 2, lane 9: eGFP plasmid 3, lane 10: eGFP plasmid 4, lane 11: eGFP plasmid 5).

**SI2. Sequencing results**

Sanger sequencing was performed using Source BioScience, using the primer TCACAAGTATGCTTGGTC (MT 57.5°C). Primers were supplied to the sequencing company at 5 pmol/µL in 5 µL of nuclease free water. Nanopore sequencing was performed using the same company. For both sequencing preparations, plasmids were prepared at 100 ng/µL in nuclease free water.

### Sanger sequencing

The raw sequences obtained from Sanger sequencing for AmpR plasmids 1, 2, 4, and 5 are provided in table S2. The text highlighted in yellow indicates the insertion sequence correctly read.

**Table S2. AmpR sequencing results from Sanger sequencing**

| Sample | Sequence |
|---|---|
| Plasmid 1 | NNNNNNNNNNNNNNNNTNNNNNNNNNNNNTGCGATGCGGCCTTGACGGCCTTCCGCCAATTCGCCCTATAGTGAGTCGTATTACGTC<br>GCGCTCACTGGCCGTCGTTTTACAACGTCGTGACTGGGAAAACCCTGGCGTTACCCAACTTAATCGCCTTGCAGCACATCCCCCTTTCG<br>CCAGCTGGCGTAATAGCGAAGAGGCCCGCACCGAAACGCCCTTCCCAACAGTTGCGCAGCCTGAATGGCGAATGGGAGCGCCCTGTA<br>GCGGCCACTCAACCCTATCTCGGTCTATTCTTTTGATTTATAAGGGATTTTGCCGATTTCGGCCTATTGGTTAAAAAATGAGCTGATTTAA<br>CAAAAATTTAACGCGAATTTTAACAAAATATTAACGCTTACAATTTAGGTGGCACTTTTCGGGGAAATGTGCGCGGAACCCCTATTTGTT<br>TATTTTTCTAAATACATTCAAATATGTATCCGCTCATGAGACAATAACCCTGATAAATGCTTCAATAATATTGAAAAAGGAAGAGTATGAGT<br>ATTCAACATTTCCGTGTCGCCCTTATTCCCTTTTTTGCGGCATTTTGCCTTCCTGTTTTTGCTCACCCAGAAACGCTGGTGAAAGTAAAAG<br>ATGCTGAAGATCAGTTGGGTGCACGAGTGGGTTACATCGAACTGGATCTCAACAGCGGTAAGATCCTTGAGAGTTTTCGCCCCGAAGA<br>ACGTTTTCCAATGATGAGCACTTTTAAAGTTCTGCTATGTGGCGCGGTATTATCCCGTATTGACGCCGGGCAAGAGCAACTCGGTCGCC<br>GCATACACTATTCTCAGAATGACTTGGTTGAGTACTCACCAGTCACAGAAAAGCATCTTACGGATGGCATGACAGTAAGAGAATTATGCA<br>GTGCTGCCATAACCATGAGTGATAACACTGCGGCCAACTTACTTCTGACAACGATCGGAGGACCGAAGGAGCTAACCGCTTTTTTGCAC<br>AACATGGGGGATCATGTAACTCGCCTTGATCGTTGGGAACCGGAGCTGAATGAAGCCATACCAAACGACGAGCGTGACACCACGATGC<br>CTGTAGCAATGGCAACAACGTTGCGCAAACTATTAACTGGCGAACTACTTACTCTAGCTTCCCGGCAACAATTAATAGACTGGATGGAG<br>GCGGATAAAGTTGCAGGACCACTTCTGNGCTCGGCCCTTCCGGCTGGCTGGTTTATTGCTGATAANNCTGAAGCCGGTGAGCGGGGT<br>TCTCCGGNNNTCATTGCANNACTGGGGCCAAATGGGAANCCCTCCCGGATTNGNNNTTATNNACCNNACGGGNNAGNNGGNNNC<br>CTTNGGATGAACNNAAAAAAANNNNANNNCT |
| Plasmid 2 | NNNNNNNNNNNNNNNNNNNNNNNNNGNCTGNNTTGGNGTNNNNGCCTGNTGNNACTAAGTNNTTGNANNNNNNNNNANACGC<br>TCNNGNAGGGGNTTGACNNNTCCGAANGNTTTCACCTCGCCCGGTTCTTNCTGAAGCCCCGTGCCTGCACCCCCTCATNGGAGATG<br>NNGNAATACCCAAGAAGGCCATCTCCAATATCCCTTGCCANCAGTTGTTCANCCTGAAGGGCCAATGGGGGNGCCCTGTACCGGCCN<br>CTCCACCCTAGCTCGGCCTATTCTTTTTATTTGTNAGGGACGNNGCTNATTTTGGCCTATTGCTTAANNCTGAGNCTATTTAACANAAAT<br>TTGACGCNANTTNNAACATAATATTAACGCGGNNATTTANGNGTCACTTTTCGGGGAAATGTGCCCTCGAATCGAGTACCGGGAACCC<br>CTATTTGTTTATTTTTCNNACTACATTCAAATATGTATCCGCTCATGAGACGGTAACCCTGATAAATGCTTCCCCACTATTGAAAAAGGAA<br>GAGTATGAGTATACAACATTTCCGNGTCNGCCTTATTCCCTTTTTTGCGGCATTTTGCCTTCGTGTTTTTGCTCACCCNGAAACNNTGGT<br>GANNNTGANAGANNNTGAAAATCAGGGGGGTGCACGAGGGGGCGTACATNNTNACTGCATCTCCCATGCGTAAGATCCTGTGANA<br>GNNTTCTCCCCCCAAGACCNCTTTCCAATGATGAGCACTTTTAAAGNTGTGCTATGTAGCGCTGTNNTAATCNCCNCATTGNNGCNNN<br>CGNAGCAGCGACTCGACCCCCGNATACGCAATT |
| Plasmid 4 | NNNNNNNNNNNNNNNNNNNNNNNNNNGGNNNGCGATGCGGCCTTGACGGCCTTCCGCCAATTCGCCCTATAGTGAGTCGTATTACG<br>TCGCGCTCACTGGCCGTCGTTTTACAACGTCGTGACTGGGAAAACCCTGGCGTTACCCAACTTAATCGCCTTGCAGCACATCCCCCTTTC<br>GCCAGCTGGCGTAATAGCGAAGAGGCCCGCACCGAAACGCCCTTCCCAACAGTTGCGCAGCCTGAATGGCGAATGGGAGCGCCCTGT<br>AGCGGCCACTCAACCCTATCTCGGTCTATTCTTTTGATTTATAAGGGATTTTGCCGATTTCGGCCTATTGGTTAAAAAATGAGCTGATTTA<br>ACAAAAATTTAACGCGAATTTTAACAAAATATTAACGCTTACAATTTAGGTGGCACTTCGAATCGAGTACCGTTTCGGGGAAATGTGCGC<br>GGAACCCCTATTTGTTTATTTTTCTAAATACATTCAAATATGTATCCGCTCATGAGACAATAACCCTGATAAATGCTTCAATAATATTGAAAA<br>AGGAAGAGTATGAGTATTCAACATTTCCGTGTCGCCCTTATTCCCTTTTTTGCGGCATTTTGCCTTCCTGTTTTTGCTCACCCAGAAACGC<br>TGGTGAAAGTAAAAGATGCTGAAGATCAGTTGGGTGCACGAGTGGGTTACATCGAACTGGATCTCAACAGCGGTAAGATCCTTGAGA<br>GTTTTCGCCCCGAAGAACGTTTTCCAATGATGAGCACTTTTAAAGTTCTGCTATGTGGCGCGGTATTATCCCGTATTGACGCCGGGCAAG<br>AGCAACTCGGTCGCCGCATACACTATTCTCAGAATGACTTGGTTGAGTACTCACCAGTCACAGAAAAGCATCTTACGGATGGCATGACA<br>GTAAGAGAATTATGCAGTGCTGCCATAACCATGNGTGATAACACTGCGGCCAACTTACTTCTGACAACGATCGGAGGANCGAANGAGC<br>TAACCGCTTTTTTGCACNNCATGGGNGGATCNTGTNACCTCCCCTTGATCGCTCNGAACCGNNACCGACGCACCCNNNCCANACCNC<br>NNCGGNACNCCNNNATCCNTG |
| Plasmid 5 | NNNNNNNNNNNNNNNNNNNNNNNNNNNNNCTGCGATGCGGCCTTGACGGCCTTCCGCCAATTCGCCCTATAGTGAGTCGTATTACGT<br>CGCGCTCACTGGCCGTCGTTTTACAACGTCGTGACTGGGAAAACCCTGGCGTTACCCAACTTAATCGCCTTGCAGCACATCCCCCTTTC<br>GCCAGCTGGCGTAATAGCGAAGAGGCCCGCACCGAAACGCCCTTCCCAACAGTTGCGCAGCCTGAATGGCGAATGGGAGCGCCCTGT<br>AGCGGCCACTCAACCCTATCTCGGTCTATTCTTTTGATTTATAAGGGATTTTGCCGATTTNANCCTATTGGTTAGAAAAAGTGCTGACTTA<br>ACAAAAATGTAANGNNAAATGTAACAAAATATTCCNAACNNACNNNNTCGGNNNNNN |

### Nanopore sequencing

The raw sequences obtained from Oxford Nanopore sequencing are provided in table S3. Red text indicates the AmpR gene, orange for the AmpR promoter, and the text highlighted in yellow shows the newly added insertion sequence. The blue highlight with the green dotted line in the sequence for AmpR plasmid 5 indicates the point at which the insertion, and part of the promoter sequence, is missing before returning to the original sequence.

**Table S3. AmpR sequencing results from nanopore sequencing**

| Sample | Sequence |
|---|---|
| Plasmid 1 | ATGAGTATTCAACATTTCCGTGTCGCCCTTATTCCCTTTTTTGCGGCATTTTGCCTTCCTGTTTTTGCTCACCCAGAAACGCTGGTGAAAG<br>TAAAAGATGCTGAAGATCAGTTGGGTGCACGAGTGGGTTACATCGAACTGGATCTCAACAGCGGTAAGATCCTTGAGAGTTTTCGCCC<br>CGAAGAACGTTTTCCAATGATGAGCACTTTTAAAGTTCTGCTATGTGGCGCGGTATTATCCCGTATTGACGCCGGGCAAGAGCAACTCG<br>GTCGCCGCATACACTATTCTCAGAATGACTTGGTTGAGTACTCACCAGTCACAGAAAAGCATCTTACGGATGGCATGACAGTAAGAGAA<br>TTATGCAGTGCTGCCATAACCATGAGTGATAACACTGCGGCCAACTTACTTCTGACAACGATCGGAGGACCGAAGGAGCTAACCGCTTT<br>TTTGCACAACATGGGGGATCATGTAACTCGCCTTGATCGTTGGGAACCGGAGCTGAATGAAGCCATACCAAACGACGAGCGTGACACC<br>ACGATGCCTGTAGCAATGGCAACAACGTTGCGCAAACTATTAACTGGCGAACTACTTACTCTAGCTTCCCGGCAACAATTAATAGACTGG<br>ATGGAGGCGGATAAAGTTGCAGGACCACTTCTGCGCTCGGCCCTTCCGGCTGGCTGGTTTATTGCTGATAAATCTGGAGCCGGTGAGC<br>GTGGTTCTCGCGGTATCATTGCAGCACTGGGGCCAGATGGTAAGCCCTCCCGTATCGTAGTTATCTACACGACGGGGAGTCAGGCAACT<br>ATGGATGAACGAAATAGACAGATCGCTGAGATAGGTGCCTCACTGATTAAGCATTGGTAACTGTCAGACCAAGTTTACTCATATATACTT<br>TAGATTGATTTAAAACTTCATTTTTAATTTAAAAGGATCTAGGTGAAGATCCTTTTTGATAATCTCATGACCAAAATCCCTTAACGTGAGTT<br>TTCGTTCCACTGAGCGTCAGACCCCGTAGAAAAGATCAAAGGATCTTCTTGAGATCCTTTTTTTCTGCGCGTAATCTGCTGCTTGCAAAC<br>AAAAAAACCACCGCTACCAGCGGTGGTTTGTTTGCCGGATCAAGAGCTACCAACTCTTTTTCCGAAGGTAACTGGCTTCAGCAGAGCG<br>CAGATACCAAATACTGTTCTTCTAGTGTAGCCGTAGTTAGGCCACCACTTCAAGAACTCTGTAGCACCGCCTACATACCTCGCTCTGCTAA<br>TCCTGTTACCAGTGGCTGCTGCCAGTGGCGATAAGTCGTGTCTTACCGGGTTGGACTCAAGACGATAGTTACCGGATAAGGCGCAGCG<br>GTCGGGCTGAACGGGGGGTTCGTGCACACAGCCCAGCTTGGAGCGAACGACCTACACCGAACTGAGATACCTACAGCGTGAGCTATG<br>AGAAAGCGCCACGCTTCCCGAAGGGAGAAAGGCGGACAGGTATCCGGTAAGCGGCAGGGTCGGAACAGGAGAGCGCACGAGGGA<br>GCTTCCAGGGGGAAACGCCTGGTATCTTTATAGTCCTGTCGGGTTTCGCCACCTCTGACTTGAGCGTCGATTTTTGTGATGCTCGTCAG<br>GGGGGCGGAGCCTATGGAAAAACGCCAGCAACGCGGCCTTTTTACGGTTCCTGGCCTTTTGCTGGCCTTTTGCTCATTAGGCACCCCA<br>GGCTTTACCCGAACGACCGAGCGCAGCGAGTCAGTGAGCGAGGAAGCGGAGAGCGCCCAATACGCAAGGAAACAGCTATGACCATG<br>TTAATGCAGCTGGCACGACAGGTTTCCCGACTGGAAAGCGGGCAGTGAGCGGAAGGCCCATGAGGCCCAGATCCAGCAGTCCGACG<br>AATTACCGGAAACTTTATATACAATTAAGAAAATTTACAAATTTGAAGTTGGAACACCCTTCCCTTAGGGCGATTTAGCCACCGCGAGCT<br>GTGTTCCGGAATTCCCGTGAAAGGGGTATTTTACACTGGCTGCGTTGCCGACTTTCTCATACTCGCGATTGCAGAGTAAAGAAATTCCC<br>ATGCTAGACCGTTGCCGGTTCCCATTGTACTTCACGGGGAGCCACAAGCGCCACAGTTCATGACCCTACGCTGAGCAGTGAATAATTAT<br>ACCATGGATTAAGCGACCGCCGTGTGCATTTCTCGCGAAACCGTGCTGACTCTTTGGTGTGATAATTCTACAGTCTGAAAGGTCTATTGA<br>GTGTCCTCGCTCTGGGCAACAAAGTATCATCGTCGGGTGAGGTTCGTAGATTTCCGGCGAGATCCTAGATACAAGGTTGCTTCAGGTCT<br>CAAGCCCGCTGATGCTAGAAAACATGTAACAGCGTGACGCACGTTATTTGTAACGTGTCGTTGCAAGAGAGTGTTCGTTGGAAGATCAT<br>GCTGGTCCGTCGTTGTTATGCGCTCTGCTGTAACCTGGCTAATGGGTTAAGGTTGTTTGAGATGAGCACACATATGGGCATCCTTCCCAA<br>TCCTGTGTCGCGCGCCTTAGCCTGAAAAGACAGGTAGGTGGCGGGATACTACACATTGTTTTCATAATGTTCCCCTTTCATATGACGGCG<br>GGGCATGAATATAGTCGTGTACCATGCCGTACGTTTATGAGATCTCGCATATCTTCACGAAGTGGGTTTCAGTCCCGATTACTCCCCAGG<br>GGTGTAGTGTACGTGATGCAATAGTAGTGGGATAGTGTCCGATGAACAACCCCAAAACCATTTTAACGGCTATCAGCGAGCACGGGCC<br>GCCAGGTCCTGGCACGAAGGATTGCGAGACTTCCGCTCACAAGTATGCTTGGTCGTGCCAAGTGTCCTCGTCTACGAGCACGGTCCTG<br>CGATGCGGCCTTGACGGCCTTCCGCCAATTCGCCCTATAGTGAGTCGTATTACGTCGCGCTCACTGGCCGTCGTTTTACAACGTCGTGAC<br>TGGGAAAACCCTGGCGTTACCCAACTTAATCGCCTTGCAGCACATCCCCCTTTCGCCAGCTGGCGTAATAGCGAAGAGGCCCGCACCG<br>AAACGCCCTTCCCAACAGTTGCGCAGCCTGAATGGCGAATGGGAGCGCCCTGTAGCGGCCACTCAACCCTATCTCGGTCTATTCTTTTG<br>ATTTATAAGGGATTTTGCCGATTTCGGCCTATTGGTTAAAAAATGAGCTGATTTAACAAAAATTTAACGCGAATTTTAACAAAATATTAAC<br>GCTTACAATTTAGGTGGCACTTTTCGGGGAAATGTGCGCGGAACCCCTATTTGTTTATTTTTCTAAATACATTCAAATATGTATCCGCTCAT<br>GAGACAATAACCCTGATAAATGCTTCAATAATATTGAAAAAGGAAGAGT |
| Plasmid 2 | ATGAGTATTCAACATTTCCGTGTCGCCCTTATTCCCTTTTTTGCGGCATTTTGCCTTCCTGTTTTTGCTCACCCAGAAACGCTGGTGAAAG<br>TAAAAGATGCTGAAGATCAGTTGGGTGCACGAGTGGGTTACATCGAACTGGATCTCAACAGCGGTAAGATCCTTGAGAGTTTTCGCCC<br>CGAAGAACGTTTTCCAATGATGAGCACTTTTAAAGTTCTGCTATGTGGCGCGGTATTATCCCGTATTGACGCCGGGCAAGAGCAACTCG<br>GTCGCCGCATACACTATTCTCAGAATGACTTGGTTGAGTACTCACCAGTCACAGAAAAGCATCTTACGGATGGCATGACAGTAAGAGAA<br>TTATGCAGTGCTGCCATAACCATGAGTGATAACACTGCGGCCAACTTACTTCTGACAACGATCGGAGGACCGAAGGAGCTAACCGCTTT<br>TTTGCACAACATGGGGGATCATGTAACTCGCCTTGATCGTTGGGAACCGGAGCTGAATGAAGCCATACCAAACGACGAGCGTGACACC<br>ACGATGCCTGTAGCAATGGCAACAACGTTGCGCAAACTATTAACTGGCGAACTACTTACTCTAGCTTCCCGGCAACAATTAATAGACTGG<br>ATGGAGGCGGATAAAGTTGCAGGACCACTTCTGCGCTCGGCCCTTCCGGCTGGCTGGTTTATTGCTGATAAATCTGGAGCCGGTGAGC<br>GTGGTTCTCGCGGTATCATTGCAGCACTGGGGCCAGATGGTAAGCCCTCCCGTATCGTAGTTATCTACACGACGGGGAGTCAGGCAACT<br>ATGGATGAACGAAATAGACAGATCGCTGAGATAGGTGCCTCACTGATTAAGCATTGGTAACTGTCAGACCAAGTTTACTCATATATACTT<br>TAGATTGATTTAAAACTTCATTTTTAATTTAAAAGGATCTAGGTGAAGATCCTTTTTGATAATCTCATGACCAAAATCCCTTAACGTGAGTT |

| | |
|---|---|
| | TTCGTTCCACTGAGCGTCAGACCCCGTAGAAAAGATCAAAGGATCTTCTTGAGATCCTTTTTTTCTGCGCGTAATCTGCTGCTTGCAAAC<br>AAAAAAACCACCGCTACCAGCGGTGGTTTGTTTGCCGGATCAAGAGCTACCAACTCTTTTTCTATACCATGGATTAAGCGACCGCCGTG<br>TGCATTTCTCGCGAAACCGTGCTGACTCTTTGGTGTGATAATTCTACAGTCTGAAAGGTCTATTGAGTGTCCTCGCTCTGGGCAACAAAG<br>TATCATCGTCGGGTGAGGTTCGTAGATTTCCGGCGAGATCCTAGATACAAGGTTGCTTCAGGTCTCAAGCCCGCTGATGCTAGAAAACA<br>TGTAACAGCGTGACGCACGTTATTTGTAACGTGTCGTTGCAAGAGAGTGTTCGTTGGAAGATCATGCTGGTCCGTCGTTGTTATGCGCT<br>CTGCTGTAACCTGGCTAATGGGTTAAGGTTGTTTGAGATGAGCACACATATGGGCATCCTTCCCAATCCTGTGTCGCGCGCCTTAGCCTG<br>AAAAGACAGGTAGGTGGCGGGATACTACACATTGTTTTCATAATGTTCCCCTTTCATATGACGGCGGGGCATGAATATAGTCGTGTACCA<br>TGCCGTACGTTTATGAGATCTCGCATATCTTCACGAAGTGGGTTTCAGTCCCGATTACTCCCCAGGGGTGTAGTGTACGTGATGCAATAG<br>TAGTGGGATAGTGTCCGATGAACAACCCCAAAACCATTTTAACGGCTATCAGCGAGCACGGGCCGCCAGGTCCTGGCACGAAGGATTG<br>CGAGACTTCCGCTCACAAGTATGCTTGGTCGTGCCAAGTGTCCTCGTCTACGAGCACGGTCCTGCGATGCGGCCTTGACGGCCTTCCGC<br>CAATTCGCCCTATAGTGAGTCGTATTACGTCGCGCTCACTGGCCGTCGTTTTACAACGTCGTGACTGGGAAAACCCTGGCGTTACCCAAC<br>TTAATCGCCTTGCAGCACATCCCCCTTTCGCCAGCTGGCGTAATAGCGAAGAGGCCCGCACCGAAACGCCCTTCCCAACAGTTGCGCA<br>GCCTGAATGGCGAATGGGAGCGCCCTGTAGCGGCCACTCAACCCTATCTCGGTCTATTCTTTTGATTTATAAGGGATTTTGCCGATTTCG<br>GCCTATTGGTTAAAAAATGAGCTGATTTAACAAAAATTTAACGCGAATTTTAACAAAATATTAACGCTTACAATTTAGGTGGCACTTCGA<br>ATCGAGTACCGCGCGGAACCCCTATTTGTTTATTTTTCTAAATACATTCAAATATGTATCCGCTCATGAGACAATAACCCTGATAAATGCTT<br>CAATAATATTGAAAAAGGAAGAGT |
| Plasmid 4 | ATGAGTATTCAACATTTCCGTGTCGCCCTTATTCCCTTTTTTGCGGCATTTTGCCTTCCTGTTTTTGCTCACCCAGAAACGCTGGTGAAAG<br>TAAAAGATGCTGAAGATCAGTTGGGTGCACGAGTGGGTTACATCGAACTGGATCTCAACAGCGGTAAGATCCTTGAGAGTTTTCGCCC<br>CGAAGAACGTTTTCCAATGATGAGCACTTTTAAAGTTCTGCTATGTGGCGCGGTATTATCCCGTATTGACGCCGGGCAAGAGCAACTCG<br>GTCGCCGCATACACTATTCTCAGAATGACTTGGTTGAGTACTCACCAGTCACAGAAAAGCATCTTACGGATGGCATGACAGTAAGAGAA<br>TTATGCAGTGCTGCCATAACCATGAGTGATAACACTGCGGCCAACTTACTTCTGACAACGATCGGAGGACCGAAGGAGCTAACCGCTTT<br>TTTGCACAACATGGGGGATCATGTAACTCGCCTTGATCGTTGGGAACCGGAGCTGAATGAAGCCATACCAAACGACGAGCGTGACACC<br>ACGATGCCTGTAGCAATGGCAACAACGTTGCGCAAACTATTAACTGGCGAACTACTTACTCTAGCTTCCCGGCAACAATTAATAGACTGG<br>ATGGAGGCGGATAAAGTTGCAGGACCACTTCTGCGCTCGGCCCTTCCGGCTGGCTGGTTTATTGCTGATAAATCTGGAGCCGGTGAGC<br>GTGGTTCTCGCGGTATCATTGCAGCACTGGGGCCAGATGGTAAGCCCTCCCGTATCGTAGTTATCTACACGACGGGGAGTCAGGCAACT<br>ATGGATGAACGAAATAGACAGATCGCTGAGATAGGTGCCTCACTGATTAAGCATTGGTAACTGTCAGACCAAGTTTACTCATATATACTT<br>TAGATTGATTTAAAACTTCATTTTTAATTTAAAAGGATCTAGGTGAAGATCCTTTTTGATAATCTCATGACCAAAATCCCTTAACGTGAGTT<br>TTCGTTCCACTGAGCGTCAGACCCCGTAGAAAAGATCAAAGGATCTTCTTGAGATCCTTTTTTTCTGCGCGTAATCTGCTGCTTGCAAAC<br>AAAAAAACCACCGCTACCAGCGGTGGTTTGTTTGCCGGATCAAGAGCTACCAACTCTTTTTCCGAAGGTAACTGGCTTCAGCAGAGCG<br>CAGATACCAAATACTGTTCTTCTAGTGTAGCCGTAGTTAGGCCACCACTTCAAGAACTCTGTAGCACCGCCTACATACCTCGCTCTGCTAA<br>TCCTGTTACCAGTGGCTGCTGCCAGTGGCGATAAGTCGTGTCTTACCGGGTTGGACTCAAGACGATAGTTACCGGATAAGGCGCAGCG<br>GTCGGGCTGAACGGGGGGTTCGTGCACACAATTCTACAGTCTGAAAGGTCTATTGAGTGTCCTCGCTCTGGGCAACAAAGTATCATCG<br>TCGGGTGAGGTTCGTAGATTTCCGGCGAGATCCTAGATACAAGGTTGCTTCAGGTCTCAAGCCCGCTGATGCTAGAAAACATGTAACAG<br>CGTGACGCACGTTATTTGTAACGTGTCGTTGCAAGAGAGTGTTCGTTGGAAGATCATGCTGGTCCGTCGTTGTTATGCGCTCTGCTGTAA<br>CCTGGCTAATGGGTTAAGGTTGTTTGAGATGAGCACACATATGGGCATCCTTCCCAATCCTGTGTCGCGCGCCTTAGCCTGAAAAGACA<br>GGTAGGTGGCGGGATACTACACATTGTTTTCATAATGTTCCCCTTTCATATGACGGCGGGGCATGAATATAGTCGTGTACCATGCCGTACG<br>TTTATGAGATCTCGCATATCTTCACGAAGTGGGTTTCAGTCCCGATTACTCCCCAGGGGTGTAGTGTACGTGATGCAATAGTAGTGGGAT<br>AGTGTCCGATGAACAACCCCAAAACCATTTTAACGGCTATCAGCGAGCACGGGCCGCCAGGTCCTGGCACGAAGGATTGCGAGACTTC<br>CGCTCACAAGTATGCTTGGTCGTGCCAAGTGTCCTCGTCTACGAGCACGGTCCTGCGATGCGGCCTTGACGGCCTTCCGCCAATTCGCC<br>CTATAGTGAGTCGTATTACGTCGCGCTCACTGGCCGTCGTTTTACAACGTCGTGACTGGGAAAACCCTGGCGTTACCCAACTTAATCGCC<br>TTGCAGCACATCCCCCTTTCGCCAGCTGGCGTAATAGCGAAGAGGCCCGCACCGAAACGCCCTTCCCAACAGTTGCGCAGCCTGAATG<br>GCGAATGGGAGCGCCCTGTAGCGGCCACTCAACCCTATCTCGGTCTATTCTTTTGATTTATAAGGGATTTTGCCGATTTCGGCCTATTGG<br>TTAAAAAATGAGCTGATTTAACAAAAATTTAACGCGAATTTTAACAAAATATTAACGCTTACAATTTAGGTGGCACTCGAATCGAGTACC<br>GGCGCGGAACCCCTATTTGTTTATTTTTCTAAATACATTCAAATATGTATCCGCTCATGAGACAATAACCCTGATAAATGCTTCAATAATATT<br>GAAAAAGGAAGAGT |
| Plasmid 5 | TTACCAATGCTTAATCAGTGAGGCACCTATCTCAGCGATCTGTCTATTTCGTTCATCCATAGTTGCCTGACTCCCCGTCGTGTAGATAACTA<br>CGATACGGGAGGGCTTACCATCTGGCCCCAGTGCTGCAATGATACCGCGAGAACCACGCTCACCGGCTCCAGATTTATCAGCAATAAAC<br>CAGCCAGCCGGAAGGGCCGAGCGCAGAAGTGGTCCTGCAACTTTATCCGCCTCCATCCAGTCTATTAATTGTTGCCGGGAAGCTAGAG<br>TAAGTAGTTCGCCAGTTAATAGTTTGCGCAACGTTGTTGCCATTGCTACAGGCATCGTGGTGTCACGCTCGTCGTTTGGTATGGCTTCAT<br>TCAGCTCCGGTTCCCAACGATCAAGGCGAGTTACATGATCCCCCATGTTGTGCAAAAAAGCGGTTAGCTCCTTCGGTCCTCCGATCGTT<br>GTCAGAAGTAAGTTGGCCGCAGTGTTATCACTCATGGTTATGGCAGCACTGCATAATTCTCTTACTGTCATGCCATCCGTAAGATGCTTTT<br>CTGTGACTGGTGAGTACTCAACCAAGTCATTCTGAGAATAGTGTATGCGGCGACCGAGTTGCTCTTGCCCGGCGTCAATACGGGATAAT<br>ACCGCGCCACATAGCAGAACTTTAAAAGTGCTCATCATTGGAAAACGTTCTTCGGGGCGAAAACTCTCAAGGATCTTACCGCTGTTGAG<br>ATCCAGTTCGATGTAACCCACTCGTGCACCCAACTGATCTTCAGCATCTTTTACTTTCACCAGCGTTTCTGGGTGAGCAAAAACAGGAA<br>GGCAAAATGCCGCAAAAAAGGGAATAAGGGCGACACGGAAATGTTGAATACTCATACTCTTCCTTTTTCAATATTATTGAAGCATTTATC<br>AGGGTTATTGTCTCATGAGCGGATACATATTTGAATGTATTTAGAAAAATAAACAAATAGGGGTTCCGCGCACATTTCCCCGAAAAGTG<br>CCACCTAAATTGTAAGCGTTAATATTTTGTTAAAATTCGCGTTAAATTTTTGTTAAATCAGCTCATTTTTTAACCAATAGGCCGAAATCGGC<br>AAAATCCCTTATAAATCAAAAGAATAGACCGAGATAGGGTTGAGTGGCCGCTACAGGGCGCTCCCATTCGCCATTCAGGCTGCGCAACT<br>GTTGGGAAGGGCGTTTCGGTGCGGGCCTCTTCGCTATTACGCCAGCTGGCGAAAGGGGGATGTGCTGCAAGGCGATTAAGTTGGGTA<br>ACGCCAGGGTTTTCCCAGTCACGACGTTGTAAAACGACGGCCAGTGAGCGCGACGTAATACGACTCACTATAGGGCGAATTGGCGGA<br>AGGCCGTCAAGGCCGCATCGCAGCCAGCTGCATTAACATGGTCATAGCTGTTTCCTTGCGTATTGGGCGCTCTCCGCTTCCTCGCTCACT<br>GACTCGCTGCGCTCGGTCGTTCGGGTAAAGCCTGGGGTGCCTAATGAGCAAAAGGCCAGCAAAAGGCCAGGAACCGTAAAAAGGCC<br>GCGTTGCTGGCGTTTTTCCATAGGCTCCGCCCCCCTGACGAGCATCACAAAAATCGACGCTCAAGTCAGAGGTGGCGAAACCCGACAG<br>GACTATAAAGATACCAGGCGTTTCCCCCTGGAAGCTCCCTCGTGCGCTCTCCTGTTCCGACCCTGCCGCTTACCGGATACCTGTCCGCCT<br>TTCTCCCTTCGGGAAGCGTGGCGCTTTCTCATAGCTCACGCTGTAGGTATCTCAGTTCGGTGTAGGTCGTTCGCTCCAAGCTGGGCTGT<br>GTGCACGAACCCCCCGTTCAGCCCGACCGCTGCGCCTTATCCGGTAACTATCGTCTTGAGTCCAACCCGGTAAGACACGACTTATCGCC<br>ACTGGCAGCAGCCACTGGTAACAGGATTAGCAGAGCGAGGTATGTAGGCGGTGCTACAGAGTTCTTGAAGTGGTGGCCTAACTACGG<br>CTACACTAGAAGAACAGTATTTGGTATCTGCGCTCTGCTGAAGCCAGTTACCTTCGGAAAAAGAGTTGGTAGCTCTTGATCCGGCAAAC<br>AAACCACCGCTGGTAGCGGTGGTTTTTTTGTTTGCAAGCAGCAGATTACGCGCAGAAAAAAAGGATCTCAAGAAGATCCTTTGATCTT<br>TTCTACGGGGTCTGACGCTCAGTGGAACGAAAACTCACGTTAAGGGATTTTGGTCATGAGATTATCAAAAAGGATCTTCACCTAGATCC<br>TTTTAAATTAAAAATGAAGTTTTAAATCAATCTAAAGTATATATGAGTAAACTTGGTCTGACAG |

### Dual gene containing plasmid characterisation

To determine the after-effect of the modification on the pCSDEST2-EGFP plasmid *in vivo*, BL21 cells were grown in liquid LB broth, and plasmids extracted using a Miniprep kit (Qiagen). The wash solutions were collected for further analysis to determine if there was any genomic integration of the plasmid DNA due to the disruption caused by the modification. Both plasmid and remaining DNA were subjected to PCR amplification (Q5 Hot Start, NEB, 98°C denaturing-10 seconds, 52°C annealing- 30 seconds, 72°C extension- 30 seconds, 30 cycles) for eGFP and AmpR, and the resulting products were run on a gel, as shown in figure S4. The primer sequences are shown below. As we discuss in the main paper, the AmpR gene is active and the eGFP gene is not, therefore primers were designed to avoid the promoter regions to determine if the genes were still present or had been excised.

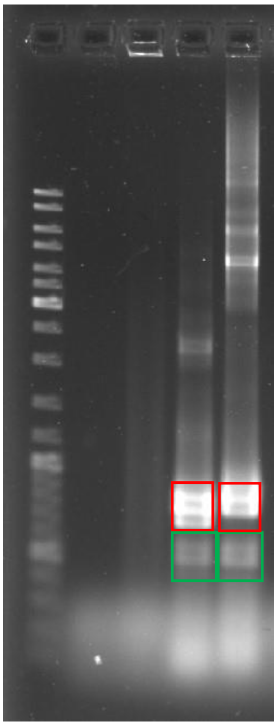

Figure S4. 1% agarose gel electrophoresis results of PCR amplified AmpR (red) and eGFP (green) genes from extracted, unmodified and modified, eGFP plasmids (lane 1: gene ruler (1kbp plus, Thermofisher), lane 2: genomic DNA, modified plasmid, lane 3: genomic DNA, unmodified plasmid, lane 4: modified plasmid amplification, lane:5 unmodified plasmid amplification).

AmpR primers

5’- TTA CCA ATG CTT AAT CAG TGA GGC ACC -3’

5’- CAC CCA GAA ACG CTG GTG AAA -3’

eGFP primers

5’- ATG GTG AGC AAG GGC GAG GA -3’

5’- CTT GTA CAG CTC GTC CAT GCC -3’

From the results of the amplification, it is possible to conclude that there is no genomic integration due to the lack of bands. The bands generated from the PCR amplified, post-growth, extracted plasmids, show that both AmpR and eGFP genes are present. Interestingly, this also showed that in the unmodified plasmids, higher weight bands than the modified plasmid were observed, indicative of restriction. The decreased yield from plasmid extraction, without the colony numbers being affected, were; modified plasmids at 25ng/ µL, and unmodified plasmids at 251ng/ µL, also indicates a decrease in copy number. In combination, these results suggested removal or a portion of the DNA containing the modified site. To determine the effect of the modification during the cell cycle, the extracted plasmid was sequenced using ONT, the results of which are shown in table S4. Red text indicates the AmpR gene, green for the eGFP promoter, and the text highlighted in purple shows the sequence missing from the modified post extraction sample.

| Sample | Sequence |
|---|---|
| Unmodified post-growth eGFP plasmid | GCCATTCAGGCTGCGCAACTGTTGGGAAGGGCGATCGGTGCGGGCCTCTTCGCTATTACGCCAGTCGACCGCCAATTCAAT ATGGCGTATATGGACTCATGCCAATTCAATATGGTGGATCTGGACCTGTGCCAATTCAATATGGCGTATATGGACTCGTGCCAA TTCAATATGGTGGATCTGGACCCCAGCCAATTCAATATGGCGGACTTGGCACCATGCCAATTCAATATGGCGGACCTGGCAC TGTGCCAACTGGGGAGGGGTCTACTTGGCACGGTGCCAAGTTTGAGGAGGGGTCTTGGCCCTGTGCCAAGTCCGCCATAT TGAATTGGCATGGTGCCAATAATGGCGGCCATATTGGCTATATGCCAGGATCAATATATAGGCAATATCCAATATGGCCCTATG CCAATATGGCTATTGGCCAGGTTCAATACTATGTATTGGCCCTATGCCATATAGTATTCCATATATGGGTTTTCCTATTGACGTAG ATAGCCCCTCCCAATGGGCGGTCCCATATACCATATATGGGGCTTCCTAATACCGCCCATAGCCACTCCCCCATTGACGTCAAT GGTCTCTATATATGGTCTTTCCTATTGACGTCATATGGGCGGTCCTATTGACGTATATGGCGCCTCCCCCATTGACGTCAATTAC GGTAAATGGCCCGCCTGGCTCAATGCCCATTGACGTCAATAGGACCACCCACCATTGACGTCAATGGGATGGCTCATTGCCC ATTCATATCCGTTCTCACGCCCCCTATTGACGTCAATGACGGTAAATGGCCCACTTGGCAGTACATCAATATCTATTAATAGTAA CTTGGCAAGTACATTACTATTGGAAGTACGCCAGGGTACATTGGCAGTACTCCCATTGACGTCAATGGCGGTAAATGGCCCG CGATGGCTGCCAAGTACATCCCCATTGACGTCAATGGGGAGGGGCAATGACGCAAATGGGCGTTCCATTGACGTAAATGGG CGGTAGGCGTGCCTAATGGGAGGTCTATATAAGCAATGCTCGTTTAGGGAACCGCCATTCTGCCTGGGGACGTCGGAGCAA |

| | |
|---|---|
| | GCTTGATTTAGGTGACACTATAGAATACAAGCTACTTGTTCTTTTTGCAGGATCCCATCGATTCGAATTCAAGGATCAACAAG<br>TTTGTACAAAAAAGCAGGCTGCCACCATGGTGAGCAAGGGCGAGGAGCTGTTCACCGGGGTGGTGCCCATCCTGGTCGA<br>GCTGGACGGCGACGTAAACGGCCACAAGTTCAGCGTGTCCGGCGAGGGCGAGGGCGATGCCACCTACGGCAAGCTGACC<br>CTGAAGTTCATCTGCACCACCGGCAAGCTGCCCGTGCCCTGGCCCACCCTCGTGACCACCCTGACCTACGGCGTGCAGTGC<br>TTCAGCCGCTACCCCGACCACATGAAGCAGCACGACTTCTTCAAGTCCGCCATGCCCGAAGGCTACGTCCAGGAGCGCACC<br>ATCTTCTTCAAGGACGACGGCAACTACAAGACCCGCGCCGAGGTGAAGTTCGAGGGCGACACCCTGGTGAACCGCATCGA<br>GCTGAAGGGCATCGACTTCAAGGAGGACGGCAACATCCTGGGGCACAAGCTGGAGTACAACTACAACAGCCACAACGTCT<br>ATATCATGGCCGACAAGCAGAAGAACGGCATCAAGGTGAACTTCAAGATCCGCCACAACATCGAGGACGGCAGCGTGCAG<br>CTCGCCGACCACTACCAGCAGAACACCCCCATCGGCGACGGCCCCGTGCTGCTGCCCGACAACCACTACCTGAGCACCCAG<br>TCCGCCCTGAGCAAAGACCCCAACGAGAAGCGCGATCACATGGTCCTGCTGGAGTTCGTGACCGCCGCCGGGATCACTCT<br>CGGCATGGACGAGCTGTACAAGCACCCAGCTTTCTTGTACAAAGTGGTGGCGAAAACGTACGACTATCTGTTCAAACTGCT<br>GCTCATCGGAGACAGCGGTGTGGGCAAGACCTGTCTGCTGTTTCGATTCAGCGAGGACGCCTTCAACACCACTTTCATCTC<br>CACCATAGGAATTGACTTCAAAATCAGAACAATTGAGCTGGATGGGAAAAAGATCAAGCTTCAAATCTGGGATACGGCGGG<br>ACAGGAGAGGTTCAGAACCATCACCACTGCATATTACAGAGGAGCAATGGGAATTATGCTGGTGTATGATATCACAAATGAG<br>AAATCATTTGACAACATCAAGAACTGGATCAGGAACATTGAGGAGCATGCATCGTCAGATGTGGAGCGAATGATATTGGGT<br>AACAGGTGTGACATGAACGACAAGAGACAAGTTTCAAAAGAAAGAGGAGAGAAGCTAGCTATTGATTACGGAATCAAGTT<br>CTTGGAAACCAGTGCAAAATCTAGCACAAATGTTGAAGAGGCATTTGTCACTCTTGCTAGAGACATCATGACGAGATTAAAC<br>AGGAAAATGAATGAGAATAACCCTTCAGGGGGAGGAGGAGGAGGAGCAGTAAAAATCACAGAGAGTCGATCCAAGAAG<br>CCAAGCTTCTTCCGCTGCATTCTGCTCTAACCCAGCCAACTTTATTATACATAGTTGCACTCGAGCCTCTAGAACTATAGTGAG<br>TCGTATTACGTAGATCCAGACATGATAAGATACATTGATGAGTTTGGACAAACCACAACTAGAATGCAGTGAAAAAAATGCT<br>TTATTTGTGAAATTTGTGATGCTATTGCTTTATTTGTAACCATTATAAGCTGCAATAAACAAGTTAACAACAACAATTGCATTCA<br>TTTTATGTTTCAGGTTCAGGGGGAGGTGTGGGAGGTTTTTTAATTCGCGGCCGCGGCGCCAATGCATTGGGCCCGGTACCC<br>AGCTTTTGTTCCCTTTAGTGAGGGTTAATTGCGCGCTTGGCGTAATCATGGTCATAGCTGTTTCCTGTGTGAAATTGTTATCC<br>GCTCACAATTCCACACAACATACGAGCCGGGAGCATAAAGTGTAAAGCCTGGGGTGCCTAATGAGTGAGCTAACTCACATT<br>AATTGCGTTGCGCTCACTGCCCGCTTTCCAGTCGGGAAACCTGTCGTGCCAGCTGCATTAATGAATCGGCCAACGCGCGGG<br>GAGAGGCGGTTTGCGTATTGGGCGCTCTTCCGCTTCCTCGCTCACTGACTCGCTGCGCTCGGTCGTTCGGCTGCGGCGAGC<br>GGTATCAGCTCACTCAAAGGCGGTAATACGGTTATCCACAGAATCAGGGGATAACGCAGGAAAGAACATGTGAGCAAAAG<br>GCCAGCAAAAGGCCAGGAACCGTAAAAAGGCCGCGTTGCTGGCGTTTTTCCATAGGCTCCGCCCCCCTGACGAGCATCAC<br>AAAAATCGACGCTCAAGTCAGAGGTGGCGAAACCCGACAGGACTATAAAGATACCAGGCGTTTCCCCCTGGAAGCTCCCTC<br>GTGCGCTCTCCTGTTCCGACCCTGCCGCTTACCGGATACCTGTCCGCCTTTCTCCCTTCGGGAAGCGTGGCGCTTTCTCATAG<br>CTCACGCTGTAGGTATCTCAGTTCGGTGTAGGTCGTTCGCTCCAAGCTGGGCTGTGTGCACGAACCCCCCGTTCAGCCCGAC<br>CGCTGCGCCTTATCCGGTAACTATCGTCTTGAGTCCAACCCGGTAAGACACGACTTATCGCCACTGGCAGCAGCCACTGGTA<br>ACAGGATTAGCAGAGCGAGGTATGTAGGCGGTGCTACAGAGTTCTTGAAGTGGTGGCCTAACTACGGCTACACTAGAAGA<br>ACAGTATTTGGTATCTGCGCTCTGCTGAAGCCAGTTACCTTCGGAAAAAGAGTTGGTAGCTCTTGATCCGGCAAACAAACCA<br>CCGCTGGTAGCGGTGGTTTTTTTGTTTGCAAGCAGCAGATTACGCGCAGAAAAAAAGGATCTCAAGAAGATCCTTTGATCT<br>TTTCTACGGGGTCTGACGCTCAGTGGAACGAAAACTCACGTTAAGGGATTTTGGTCATGAGATTATCAAAAAGGATCTTCAC<br>CTAGATCCTTTTAAATTAAAAATGAAGTTTTAAATCAATCTAAAGTATATATGAGTAAACTTGGTCTGACAGTTACCAATGCTTA<br>ATCAGTGAGGCACCTATCTCAGCGATCTGTCTATTTCGTTCATCCATAGTTGCCTGACTCCCCGTCGTGTAGATAACTACGATA<br>CGGGAGGGCTTACCATCTGGCCCCAGTGCTGCAATGATACCGCGAGACCCACGCTCACCGGCTCCAGATTTATCAGCAATAA<br>ACCAGCCAGCCGGAAGGGCCGAGCGCAGAAGTGGTCCTGCAACTTTATCCGCCTCCATCCAGTCTATTAATTGTTGCCGGG<br>AAGCTAGAGTAAGTAGTTCGCCAGTTAATAGTTTGCGCAACGTTGTTGCCATTGCTACAGGCATCGTGGTGTCACGCTCGTC<br>GTTTGGTATGGCTTCATTCAGCTCCGGTTCCCAACGATCAAGGCGAGTTACATGATCCCCCATGTTGTGCAAAAAAGCGGTT<br>AGCTCCTTCGGTCCTCCGATCGTTGTCAGAAGTAAGTTGGCCGCAGTGTTATCACTCATGGTTATGGCAGCACTGCATAATTC<br>TCTTACTGTCATGCCATCCGTAAGATGCTTTTCTGTGACTGGTGAGTACTCAACCAAGTCATTCTGAGAATAGTGTATGCGGC<br>GACCGAGTTGCTCTTGCCCGGCGTCAATACGGGATAATACCGCGCCACATAGCAGAACTTTAAAAGTGCTCATCATTGGAAA<br>ACGTTCTTCGGGGCGAAAACTCTCAAGGATCTTACCGCTGTTGAGATCCAGTTCGATGTAACCCACTCGTGCACCCAACTGA<br>TCTTCAGCATCTTTTACTTTCACCAGCGTTTCTGGGTGAGCAAAAACAGGAAGGCAAAATGCCGCAAAAAAGGGAATAAG<br>GGCGACACGGAAATGTTGAATACTCATACTCTTCCTTTTTCAATATTATTGAAGCATTTATCAGGGTTATTGTCTCATGAGCGG<br>ATACATATTTGAATGTATTTAGAAAAATAAACAAATAGGGGTTCCGCGCACATTTCCCCGAAAAGTGCCACCTAAATTGTAAG<br>CGTTAATATTTTGTTAAAATTCGCGTTAAATTTTTGTTAAATCAGCTCATTTTTTAACCAATAGGCCGAAATCGGCAAAATCCC<br>TTATAAATCAAAAGAATAGACCGAGATAGGGTTGAGTGTTGTTCCAGTTTGGAACAAGAGTCCACTATTAAAGAACGTGGA<br>CTCCAACGTCAAAGGGCGAAAAACCGTCTATCAGGGCGATGGCCCACTACGTGAACCATCACCCTAATCAAGTTTTTTGGG<br>GTCGAGGTGCCGTAAAGCACTAAATCGGAACCCTAAAGGGAGCCCCCGATTTAGAGCTTGACGGGGAAAGCCGGCGAAC<br>GTGGCGAGAAAGGAAGGGAAGAAAGCGAAAGGAGCGGGCGCTAGGGCGCTGGCAAGTGTAGCGGTCACGCTGCGCGT<br>AACCACCACACCCGCCGCGCTTAATGCGCCGCTACAGGGCGCGTCCCATTC |
| Modified post-growth eGFP plasmid | ACGCGCCCTGTAGCGGCGCATTAAGCGCGGCGGGTGTGGTGGTTACGCGCAGCGTGACCGCTACACTTGCCAGCGCCCTA<br>GCGCCCGCTCCTTTCGCTTTCTTCCCTTCCTTTCTCGCCACGTTCGCCGGCTTTCCCCGTCAAGCTCTAAATCGGGGGCTCCC<br>TTTAGGGTTCCGATTTAGTGCTTTACGGCACCTCGACCCCAAAAAACTTGATTAGGGTGATGGTTCACGTAGTGGGCCATCG<br>CCCTGATAGACGGTTTTTCGCCCTTTGACGTTGGAGTCCACGTTCTTTAATAGTGGACTCTTGTTCCAAACTGGAACAACAC<br>TCAACCCTATCTCGGTCTATTCTTTTGATTTATAAGGGATTTTGCCGATTTCGGCCTATTGGTTAAAAAATGAGCTGATTTAAC<br>AAAAATTTAACGCGAATTTTAACAAAATATTAACGCTTACAATTTAGGTGGCACTTTTCGGGGAAATGTGCGCGGAACCCCT<br>ATTTGTTTATTTTTCTAAATACATTCAAATATGTATCCGCTCATGAGACAATAACCCTGATAAATGCTTCAATAATATTGAAAAAG<br>GAAGAGTATGAGTATTCAACATTTCCGTGTCGCCCTTATTCCCTTTTTTGCGGCATTTTGCCTTCCTGTTTTTGCTCACCCAGA |

| | |
|---|---|
| | AACGCTGGTGAAAGTAAAAGATGCTGAAGATCAGTTGGGTGCACGAGTGGGTTACATCGAACTGGATCTCAACAGCGGTA<br>AGATCCTTGAGAGTTTTCGCCCCGAAGAACGTTTTCCAATGATGAGCACTTTTAAAGTTCTGCTATGTGGCGCGGTATTATCC<br>CGTATTGACGCCGGGCAAGAGCAACTCGGTCGCCGCATACACTATTCTCAGAATGACTTGGTTGAGTACTCACCAGTCACAG<br>AAAAGCATCTTACGGATGGCATGACAGTAAGAGAATTATGCAGTGCTGCCATAACCATGAGTGATAACACTGCGGCCAACTT<br>ACTTCTGACAACGATCGGAGGACCGAAGGAGCTAACCGCTTTTTTGCACAACATGGGGGATCATGTAACTCGCCTTGATCG<br>TTGGGAACCGGAGCTGAATGAAGCCATACCAAACGACGAGCGTGACACCACGATGCCTGTAGCAATGGCAACAACGTTGC<br>GCAAACTATTAACTGGCGAACTACTTACTCTAGCTTCCCGGCAACAATTAATAGACTGGATGGAGGCGGATAAAGTTGCAGG<br>ACCACTTCTGCGCTCGGCCCTTCCGGCTGGCTGGTTTATTGCTGATAAATCTGGAGCCGGTGAGCGTGGGTCTCGCGGTATC<br>ATTGCAGCACTGGGGCCAGATGGTAAGCCCTCCCGTATCGTAGTTATCTACACGACGGGGAGTCAGGCAACTATGGATGAA<br>CGAAATAGACAGATCGCTGAGATAGGTGCCTCACTGATTAAGCATTGGTAACTGTCAGACCAAGTTTACTCATATATACTTTA<br>GATTGATTTAAAACTTCATTTTTAATTTAAAAGGATCTAGGTGAAGATCCTTTTTGATAATCTCATGACCAAAATCCCTTAACG<br>TGAGTTTTCGTTCCACTGAGCGTCAGACCCCGTAGAAAAGATCAAAGGATCTTCTTGAGATCCTTTTTTTCTGCGCGTAATCT<br>GCTGCTTGCAAACAAAAAAACCACCGCTACCAGCGGTGGTTTGTTTGCCGGATCAAGAGCTACCAACTCTTTTTCCGAAGG<br>TAACTGGCTTCAGCAGAGCGCAGATACCAAATACTGTTCTTCTAGTGTAGCCGTAGTTAGGCCACCACTTCAAGAACTCTGT<br>AGCACCGCCTACATACCTCGCTCTGCTAATCCTGTTACCAGTGGCTGCTGCCAGTGGCGATAAGTCGTGTCTTACCGGGTTG<br>GACTCAAGACGATAGTTACCGGATAAGGCGCAGCGGTCGGGCTGAACGGGGGGTTCGTGCACACAGCCCAGCTTGGAGC<br>GAACGACCTACACCGAACTGAGATACCTACAGCGTGAGCTATGAGAAAGCGCCACGCTTCCCGAAGGGAGAAAGGCGGA<br>CAGGTATCCGGTAAGCGGCAGGGTCGGAACAGGAGAGCGCACGAGGGAGCTTCCAGGGGGAAACGCCTGGTATCTTTAT<br>AGTCCTGTCGGGTTTCGCCACCTCTGACTTGAGCGTCGATTTTTGTGATGCTCGTCAGGGGGGCGGAGCCTATGGAAAAAC<br>GCCAGCAACGCGGCCTTTTTACGGTTCCTGGCCTTTTGCTGGCCTTTTGCTCACATGTTCTTTCCTGCGTTATCCCCTGATTCT<br>GTGGATAACCGTATTACCGCCTTTGAGTGAGCTGATACCGCTCGCCGCAGCCGAACGACCGAGCGCAGCGAGTCAGTGAG<br>CGAGGAAGCGGAAGAGCGCCCAATACGCAAACCGCCTCTCCCCGCGCGTTGGCCGATTCATTAATGCAGGATCTCGATCCC<br>GCGAAATTAATACGACTCACTATAGGGAGACCACAACGGTTTCCCTCTAGAAATAATTTTGTTTAACTTTAAGAAGGAGATAT<br>ACATATGCGGGGTTCTCATCATCATCATCATCATGGTATGGCTAGCATGACTGGTGGACAGCAAATGGGTCGGGATCTGTACG<br>ACGATGACGATAAGGATCCCATGGTGAGCAAGGGCGAGGAGCTGTTCACCGGGGTGGTGCCCATCCTGGTCGAGCTGGAC<br>GGCGACGTAAACGGCCACAAGTTCAGCGTGTCCGGCGAGGGCGAGGGCGATGCCACCTACGGCAAGCTGACCCTGAAGT<br>TCATCTGCACCACCGGCAAGCTGCCCGTGCCCTGGCCCACCCTCGTGACCACCCTGACCTACGGCGTGCAGTGCTTCAGCC<br>GCTACCCCGACCACATGAAGCAGCACGACTTCTTCAAGTCCGCCATGCCCGAAGGCTACGTCCAGGAGCGCACCATCTTCT<br>TCAAGGACGACGGCAACTACAAGACCCGCGCCGAGGTGAAGTTCGAGGGCGACACCCTGGTGAACCGCATCGAGCTGAA<br>GGGCATCGACTTCAAGGAGGACGGCAACATCCTGGGGCACAAGCTGGAGTACAACTACAACAGCCACAACGTCTATATCAT<br>GGCCGACAAGCAGAAGAACGGCATCAAGGTGAACTTCAAGATCCGCCACAACATCGAGGACGGCAGCGTGCAGCTCGCC<br>GACCACTACCAGCAGAACACCCCCATCGGCGACGGCCCCGTGCTGCTGCCCGACAACCACTACCTGAGCACCCAGTCCGCC<br>CTGAGCAAAGACCCCAACGAGAAGCGCGATCACATGGTCCTGCTGGAGTTCGTGACCGCCGCCGGGATCACTCTCGGCAT<br>GGACGAGCTGTACAAGTAAGAATTCGAAGCTTGATCCGGCTGCTAACAAAGCCCGAAAGGAAGCTGAGTTGGCTGCTGCC<br>ACCGCTGAGCAATAACTAGCATAACCCCTTGGGGCCTCTAAACGGGTCTTGAGGGGTTTTTTGCTGAAAGGAGGAACTATA<br>TCCGGATCTGGCGTAATAGCGAAGAGGCCCGCACCGATCGCCCTTCCCAACAGTTGCGCAGCCTGAATGGCGAATGGG |

From the results of the sequencing, we hypothesize that, during replication the bacteria employes its restriction enzymes to remove the section that it is unable to replicate, then repairs the damaged region, reforming a smaller plasmid. This would be an energy intensive process and therefore it is likely that the bacteria prioritizes the modification of fewer plasmids, ejecting or not copying excess plasmids, reducing the copy number as the cells divide, and therefore the yield obtained from extraction.

Whilst the mechanism by which this process occurs is currently unknown, we suggest that the AdoHcy azide triggers the host excision and repair pathways. This would cause the removal of the modification site and surrounding DNA then reseal the gaps left to produce a new, smaller plasmid missing vital sequences to create an active gene.

### SI3. Ligation insertion sequence stability

Initially, the insertion sequence was ligated into the plasmid directly after the initial digestion step. However, it was noticed that the colony count was significantly lower than the $CH_3$ or MTC5 modified plasmids in the ampicillin containing plates. As the sequences are identical, we thought this was unusual, therefore the ligation plasmids were re-prepared with an additional heating step at 50°C, post ligation. We found that the additional heating step assisted with stabilising the formation of the final plasmid, improving the overall transformation, and bringing the colony count to similar levels to those observed in the other modified plasmids, as shown in figure S4.

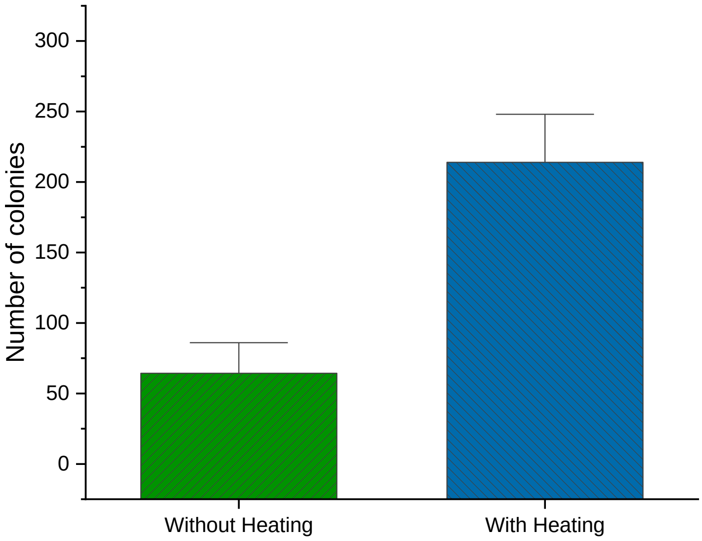


Figure S5. Bar chart showing the difference in colony count when the ligation insertion plasmid is prepared without and with a 50°C heating step respectively.

### SI4. Statistical analysis

For statistical analysis, the presence of an antibiotic was selected as one factor (two levels) and the plasmid modification as the second factor (five levels) for a two-way ANOVA comparison. Normality testing was performed using the Shapiro-Wilk test, table S4. Different plasmids were analysed individually to determine the effect of plasmid modification on bacterial growth. Results from this analysis show significant differences in the colony numbers observed for the native bacteria, bacteria with plasmids 1-4, and with plasmid 5. A summary table of the two-way ANOVA results is shown in table S5, with the individual condition statistics shown in tables S6-S9. Statistical analysis of the colony counts for the bacteria transformed with the eGFP plasmid is shown in table S10.

**Table S4. Normality testing of all bacterial conditions using the Shapiro-Wilk test**

| Bacterial condition | Plasmid condition | DF | Statistic | p-value | Decision at level(5%) |
|---|---|---|---|---|---|
| DH5α No Ampicllin | No transformation | 3 | 0.92827 | 0.48216 | Can't reject normality |
| | Control | 3 | 0.96889 | 0.66139 | Can't reject normality |
| | Ligation insert | 3 | 0.99324 | 0.84283 | Can't reject normality |
| | CH3 | 3 | 0.86502 | 0.28148 | Can't reject normality |
| | MTC5 | 3 | 0.84211 | 0.21956 | Can't reject normality |
| | AdoHcy-azide | 3 | 0.97417 | 0.6917 | Can't reject normality |
| DH5α Ampicllin | No transformation | 3 | -- | -- | a* |
| | Control | 3 | 0.98584 | 0.77223 | Can't reject normality |
| | Ligation insert | 3 | 0.99933 | 0.95064 | Can't reject normality |
| | CH3 | 3 | 0.99355 | 0.84647 | Can't reject normality |
| | MTC5 | 3 | 0.98792 | 0.78971 | Can't reject normality |
| | AdoHcy-azide | 3 | -- | -- | a* |
| BL21 No Kanamycin | No transformation | 3 | 0.88479 | 0.33861 | Can't reject normality |
| | Control | 3 | 0.99448 | 0.85798 | Can't reject normality |
| | Ligation insert | 3 | 0.99708 | 0.89673 | Can't reject normality |
| | CH3 | 3 | 0.96429 | 0.63689 | Can't reject normality |
| | MTC5 | 3 | 1 | 1 | Can't reject normality |
| | AdoHcy-azide | 3 | 0.99787 | 0.91185 | Can't reject normality |
| BL21 Kanamycin | No transformation | 3 | -- | -- | a* |
| | Control | 3 | 0.90303 | 0.39523 | Can't reject normality |
| | Ligation insert | 3 | 0.97453 | 0.6939 | Can't reject normality |
| | CH3 | 3 | 0.99817 | 0.91837 | Can't reject normality |
| | MTC5 | 3 | 0.99668 | 0.88986 | Can't reject normality |
| | AdoHcy-azide | 3 | -- | -- | a* |
| BL21 No Ampicllin | No transformation | 3 | 0.98419 | 0.75924 | Can't reject normality |
| | Control | 3 | 0.88589 | 0.34191 | Can't reject normality |
| | Ligation insert | 3 | 0.80959 | 0.13759 | Can't reject normality |
| | CH3 | 3 | 0.97959 | 0.72623 | Can't reject normality |
| | MTC5 | 3 | 0.91873 | 0.44788 | Can't reject normality |
| | AdoHcy-azide | 3 | 0.81964 | 0.16229 | Can't reject normality |
| BL21 Ampicllin | No transformation | 3 | -- | -- | a* |
| | Control | 3 | 0.96429 | 0.63689 | Can't reject normality |
| | Ligation insert | 3 | 0.98782 | 0.78881 | Can't reject normality |
| | CH3 | 3 | 0.98522 | 0.76726 | Can't reject normality |
| | MTC5 | 3 | 0.93623 | 0.51241 | Can't reject normality |
| | AdoHcy-azide | 3 | -- | -- | a* |
| DH5α No Kanamycin | No transformation | 3 | 0.9453 | 0.54913 | Can't reject normality |
| | Control | 3 | 0.89286 | 0.36311 | Can't reject normality |
| | Ligation insert | 3 | 0.99395 | 0.85129 | Can't reject normality |
| | CH3 | 3 | 0.86867 | 0.29175 | Can't reject normality |
| | MTC5 | 3 | 0.94391 | 0.54334 | Can't reject normality |
| | AdoHcy-azide | 3 | 0.99436 | 0.85645 | Can't reject normality |
| DH5α Kanamycin | No transformation | 3 | -- | -- | a* |
| | Control | 3 | 0.85465 | 0.25297 | Can't reject normality |
| | Ligation insert | 3 | 0.99976 | 0.9702 | Can't reject normality |
| | CH3 | 3 | 0.93938 | 0.52488 | Can't reject normality |
| | MTC5 | 3 | 0.92308 | 0.46326 | Can't reject normality |
| | AdoHcy-azide | 3 | -- | -- | a* |
| DH5α GFP plasmid, Amicillin plates | Control | 3 | 0.84211 | 0.21956 | Can't reject normality |
| | No transformation | 3 | -- | -- | a* |
| | AdoHcy-Azide | 3 | 0.76743 | 0.03891 | Reject normality |
| BL21 GFP plasmid, Amicillin plates | Control | 3 | 0.99973 | 0.9685 | Can't reject normality |
| | No transformation | 3 | -- | -- | a* |
| | AdoHcy-Azide | 3 | 0.78936 | 0.08934 | Can't reject normality |

a*: All colony counts were 0.

### Table S5. Summary table of the statistically significant results from the colony growth assay

| DH5α Ampicillin resistance gene | | | | | BL21 Ampicillin resistance gene | | | | |
|---|---|---|---|---|---|---|---|---|---|
| Antibiotic | Modification | Antibiotic | Modification | Prob | Antibiotic | Modification | Antibiotic | Modification | Prob |
| AMP | Control | AMP | No transformation | 3.18E-08 | AMP | No transformation | NoAMP | No transformation | 2.83E-07 |
| AMP | AdoHcy | AMP | Control | 3.18E-08 | AMP | AdoHcy | NoAMP | No transformation | 2.83E-07 |
| AMP | Ligation insert | AMP | No transformation | 3.96E-08 | AMP | No transformation | NoAMP | Control | 3.42E-06 |
| AMP | AdoHcy | AMP | Ligation insert | 3.96E-08 | AMP | AdoHcy | NoAMP | Control | 3.42E-06 |
| AMP | No transformation | NoAMP | Control | 5.15E-08 | AMP | No transformation | NoAMP | MTC5 | 5.59E-06 |
| AMP | No transformation | NoAMP | MTC5 | 5.15E-08 | AMP | AdoHcy | NoAMP | MTC5 | 5.59E-06 |
| AMP | AdoHcy | NoAMP | Control | 5.15E-08 | AMP | No transformation | NoAMP | Ligation insert | 6.18E-06 |
| AMP | AdoHcy | NoAMP | MTC5 | 5.15E-08 | AMP | No transformation | NoAMP | AdoHcy | 6.18E-06 |
| AMP | CH3 | AMP | No transformation | 5.69E-08 | AMP | AdoHcy | NoAMP | Ligation insert | 6.18E-06 |
| AMP | AdoHcy | AMP | CH3 | 5.69E-08 | AMP | AdoHcy | NoAMP | AdoHcy | 6.18E-06 |
| AMP | No transformation | NoAMP | CH3 | 1.34E-07 | AMP | Control | AMP | No transformation | 8.09E-06 |
| AMP | AdoHcy | NoAMP | CH3 | 1.34E-07 | AMP | AdoHcy | AMP | Control | 8.09E-06 |
| AMP | MTC5 | AMP | No transformation | 2.30E-07 | AMP | Ligation insert | AMP | No transformation | 8.99E-06 |
| AMP | AdoHcy | AMP | MTC5 | 2.30E-07 | AMP | AdoHcy | AMP | Ligation insert | 8.99E-06 |
| AMP | No transformation | NoAMP | No transformation | 2.66E-07 | AMP | No transformation | NoAMP | CH3 | 1.63E-05 |
| AMP | AdoHcy | NoAMP | No transformation | 2.66E-07 | AMP | AdoHcy | NoAMP | CH3 | 1.63E-05 |
| AMP | No transformation | NoAMP | AdoHcy | 4.90E-07 | AMP | CH3 | AMP | No transformation | 2.20E-05 |
| AMP | AdoHcy | NoAMP | AdoHcy | 4.90E-07 | AMP | AdoHcy | AMP | CH3 | 2.20E-05 |
| AMP | No transformation | NoAMP | Ligation insert | 1.23E-06 | AMP | MTC5 | AMP | No transformation | 3.70E-05 |
| AMP | AdoHcy | NoAMP | Ligation insert | 1.23E-06 | AMP | AdoHcy | AMP | MTC5 | 3.70E-05 |
| NoAMP | Ligation insert | NoAMP | No transformation | 0.00178 | AMP | MTC5 | NoAMP | No transformation | 0.01388 |
| NoAMP | AdoHcy | NoAMP | No transformation | 0.00469 | AMP | CH3 | NoAMP | No transformation | 0.02884 |
| AMP | MTC5 | NoAMP | No transformation | 0.01008 | NoAMP | CH3 | NoAMP | No transformation | 0.04447 |
| NoAMP | CH3 | NoAMP | No transformation | 0.01642 | | | | | |
| DH5α Kanamycin resistance gene | | | | | BL21 Kanamycin resistance gene | | | | |
| Antibiotic | Modification | Antibiotic | Modification | Prob | Antibiotic | Modification | Antibiotic | Modification | Prob |
| Kan | No transformation | NoAMP | No transformation | 4.14E-06 | Kan | No transformation | NoKan | No transformation | 4.20E-07 |
| Kan | AdoHcy | NoAMP | No transformation | 4.14E-06 | Kan | AdoHcy | NoKan | No transformation | 4.20E-07 |
| Kan | CH3 | AMP | No transformation | 6.12E-05 | Kan | Ligation insert | Kan | No transformation | 1.73E-06 |
| Kan | AdoHcy | AMP | CH3 | 6.12E-05 | Kan | AdoHcy | Kan | Ligation insert | 1.73E-06 |
| Kan | Ligation insert | AMP | No transformation | 7.34E-05 | Kan | No transformation | NoKan | Control | 1.97E-06 |
| Kan | AdoHcy | AMP | Ligation insert | 7.34E-05 | Kan | AdoHcy | NoKan | Control | 1.97E-06 |
| Kan | No transformation | NoAMP | Ligation insert | 8.11E-05 | Kan | No transformation | NoKan | MTC5 | 3.54E-06 |
| Kan | AdoHcy | NoAMP | Ligation insert | 8.11E-05 | Kan | AdoHcy | NoKan | MTC5 | 3.54E-06 |
| Kan | Control | AMP | No transformation | 8.61E-05 | Kan | No transformation | NoKan | Ligation insert | 4.11E-06 |
| Kan | AdoHcy | AMP | Control | 8.61E-05 | Kan | AdoHcy | NoKan | Ligation insert | 4.11E-06 |
| Kan | No transformation | NoAMP | Control | 9.73E-05 | Kan | CH3 | Kan | No transformation | 5.30E-06 |
| Kan | AdoHcy | NoAMP | Control | 9.73E-05 | Kan | AdoHcy | Kan | CH3 | 5.30E-06 |
| Kan | No transformation | NoAMP | AdoHcy | 1.17E-04 | Kan | Control | Kan | No transformation | 7.74E-06 |
| Kan | AdoHcy | NoAMP | AdoHcy | 1.17E-04 | Kan | AdoHcy | Kan | Control | 7.74E-06 |
| Kan | MTC5 | AMP | No transformation | 1.64E-04 | Kan | No transformation | NoKan | CH3 | 1.25E-05 |
| Kan | AdoHcy | AMP | MTC5 | 1.64E-04 | Kan | AdoHcy | NoKan | CH3 | 1.25E-05 |
| Kan | No transformation | NoAMP | CH3 | 1.75E-04 | Kan | No transformation | NoKan | AdoHcy | 1.54E-05 |
| Kan | AdoHcy | NoAMP | CH3 | 1.75E-04 | Kan | AdoHcy | NoKan | AdoHcy | 1.54E-05 |
| Kan | No transformation | NoAMP | MTC5 | 2.21E-04 | Kan | MTC5 | Kan | No transformation | 2.02E-05 |
| Kan | AdoHcy | NoAMP | MTC5 | 2.21E-04 | Kan | AdoHcy | Kan | MTC5 | 2.02E-05 |

**Table S6. Two-way ANOVA results of DH5α with and without modified plasmids on control and ampicillin plates (statistically significant results highlighted in red).**

| Factor A | Factor B | Factor A | Factor B | MeanDiff | SEM | q Value | Prob | Alpha | Sig | LCL | UCL |
|---|---|---|---|---|---|---|---|---|---|---|---|
| NoAMP | Control | NoAMP | No transformat | -65 | 20.63169 | 4.45547 | 0.1272 | 0.05 | 0 | -139.39 | 9.39027 |
| NoAMP | Ligation ins | NoAMP | No transformat | -104 | 20.63169 | 7.12875 | 0.00178 | 0.05 | 1 | -178.39 | -29.6097 |
| NoAMP | Ligation ins | NoAMP | Control | -39 | 20.63169 | 2.67328 | 0.75395 | 0.05 | 0 | -113.39 | 35.39027 |
| NoAMP | CH3 | NoAMP | No transformat | -84.6667 | 20.63169 | 5.80354 | 0.01642 | 0.05 | 1 | -159.057 | -10.2764 |
| NoAMP | CH3 | NoAMP | Control | -19.6667 | 20.63169 | 1.34807 | 0.99746 | 0.05 | 0 | -94.0569 | 54.72361 |
| NoAMP | CH3 | NoAMP | Ligation insert | 19.33333 | 20.63169 | 1.32522 | 0.99781 | 0.05 | 0 | -55.0569 | 93.72361 |
| NoAMP | MTC5 | NoAMP | No transformat | -65 | 20.63169 | 4.45547 | 0.1272 | 0.05 | 0 | -139.39 | 9.39027 |
| NoAMP | MTC5 | NoAMP | Control | 0 | 20.63169 | 0 | 1 | 0.05 | 0 | -74.3903 | 74.39027 |
| NoAMP | MTC5 | NoAMP | Ligation insert | 39 | 20.63169 | 2.67328 | 0.75395 | 0.05 | 0 | -35.3903 | 113.3903 |
| NoAMP | MTC5 | NoAMP | CH3 | 19.66667 | 20.63169 | 1.34807 | 0.99746 | 0.05 | 0 | -54.7236 | 94.05694 |
| NoAMP | AdoHcy | NoAMP | No transformat | -95.6667 | 20.63169 | 6.55754 | 0.00469 | 0.05 | 1 | -170.057 | -21.2764 |
| NoAMP | AdoHcy | NoAMP | Control | -30.6667 | 20.63169 | 2.10207 | 0.93047 | 0.05 | 0 | -105.057 | 43.72361 |
| NoAMP | AdoHcy | NoAMP | Ligation insert | 8.33333 | 20.63169 | 0.57121 | 1 | 0.05 | 0 | -66.0569 | 82.72361 |
| NoAMP | AdoHcy | NoAMP | CH3 | -11 | 20.63169 | 0.754 | 0.99999 | 0.05 | 0 | -85.3903 | 63.39027 |
| NoAMP | AdoHcy | NoAMP | MTC5 | -30.6667 | 20.63169 | 2.10207 | 0.93047 | 0.05 | 0 | -105.057 | 43.72361 |
| AMP | No transfor | NoAMP | No transformat | -272 | 20.63169 | 18.64443 | 2.66E-07 | 0.05 | 1 | -346.39 | -197.61 |
| AMP | No transfor | NoAMP | Control | -207 | 20.63169 | 14.18896 | 5.15E-08 | 0.05 | 1 | -281.39 | -132.61 |
| AMP | No transfor | NoAMP | Ligation insert | -168 | 20.63169 | 11.51568 | 1.23E-06 | 0.05 | 1 | -242.39 | -93.6097 |
| AMP | No transfor | NoAMP | CH3 | -187.333 | 20.63169 | 12.84089 | 1.34E-07 | 0.05 | 1 | -261.724 | -112.943 |
| AMP | No transfor | NoAMP | MTC5 | -207 | 20.63169 | 14.18896 | 5.15E-08 | 0.05 | 1 | -281.39 | -132.61 |
| AMP | No transfor | NoAMP | AdoHcy | -176.333 | 20.63169 | 12.08689 | 4.90E-07 | 0.05 | 1 | -250.724 | -101.943 |
| AMP | Control | NoAMP | No transformat | -49.3333 | 20.63169 | 3.38159 | 0.44713 | 0.05 | 0 | -123.724 | 25.05694 |
| AMP | Control | NoAMP | Control | 15.66667 | 20.63169 | 1.07388 | 0.99967 | 0.05 | 0 | -58.7236 | 90.05694 |
| AMP | Control | NoAMP | Ligation insert | 54.66667 | 20.63169 | 3.74716 | 0.30754 | 0.05 | 0 | -19.7236 | 129.0569 |
| AMP | Control | NoAMP | CH3 | 35.33333 | 20.63169 | 2.42195 | 0.84578 | 0.05 | 0 | -39.0569 | 109.7236 |
| AMP | Control | NoAMP | MTC5 | 15.66667 | 20.63169 | 1.07388 | 0.99967 | 0.05 | 0 | -58.7236 | 90.05694 |
| AMP | Control | NoAMP | AdoHcy | 46.33333 | 20.63169 | 3.17595 | 0.53579 | 0.05 | 0 | -28.0569 | 120.7236 |
| AMP | Control | NoAMP | No transformat | 222.6667 | 20.63169 | 15.26284 | 3.18E-08 | 0.05 | 1 | 148.2764 | 297.0569 |
| AMP | Ligation ins | NoAMP | No transformat | -58 | 20.63169 | 3.97565 | 0.23611 | 0.05 | 0 | -132.39 | 16.39027 |
| AMP | Ligation ins | NoAMP | Control | 7 | 20.63169 | 0.47982 | 1 | 0.05 | 0 | -67.3903 | 81.39027 |
| AMP | Ligation ins | NoAMP | Ligation insert | 46 | 20.63169 | 3.1531 | 0.54589 | 0.05 | 0 | -28.3903 | 120.3903 |
| AMP | Ligation ins | NoAMP | CH3 | 26.66667 | 20.63169 | 1.82789 | 0.97231 | 0.05 | 0 | -47.7236 | 101.0569 |
| AMP | Ligation ins | NoAMP | MTC5 | 7 | 20.63169 | 0.47982 | 1 | 0.05 | 0 | -67.3903 | 81.39027 |
| AMP | Ligation ins | NoAMP | AdoHcy | 37.66667 | 20.63169 | 2.58189 | 0.7895 | 0.05 | 0 | -36.7236 | 112.0569 |
| AMP | Ligation ins | AMP | No transformat | 214 | 20.63169 | 14.66878 | 3.96E-08 | 0.05 | 1 | 139.6097 | 288.3903 |
| AMP | Ligation ins | AMP | Control | -8.66667 | 20.63169 | 0.59406 | 1 | 0.05 | 0 | -83.0569 | 65.72361 |
| AMP | CH3 | NoAMP | No transformat | -67 | 20.63169 | 4.59256 | 0.10517 | 0.05 | 0 | -141.39 | 7.39027 |
| AMP | CH3 | NoAMP | Control | -2 | 20.63169 | 0.13709 | 1 | 0.05 | 0 | -76.3903 | 72.39027 |
| AMP | CH3 | NoAMP | Ligation insert | 37 | 20.63169 | 2.53619 | 0.80641 | 0.05 | 0 | -37.3903 | 111.3903 |
| AMP | CH3 | NoAMP | CH3 | 17.66667 | 20.63169 | 1.21097 | 0.99901 | 0.05 | 0 | -56.7236 | 92.05694 |
| AMP | CH3 | NoAMP | MTC5 | -2 | 20.63169 | 0.13709 | 1 | 0.05 | 0 | -76.3903 | 72.39027 |
| AMP | CH3 | NoAMP | AdoHcy | 28.66667 | 20.63169 | 1.96498 | 0.9547 | 0.05 | 0 | -45.7236 | 103.0569 |
| AMP | CH3 | AMP | No transformat | 205 | 20.63169 | 14.05187 | 5.69E-08 | 0.05 | 1 | 130.6097 | 279.3903 |
| AMP | CH3 | AMP | Control | -17.6667 | 20.63169 | 1.21097 | 0.99901 | 0.05 | 0 | -92.0569 | 56.72361 |
| AMP | CH3 | AMP | Ligation insert | -9 | 20.63169 | 0.61691 | 1 | 0.05 | 0 | -83.3903 | 65.39027 |
| AMP | MTC5 | NoAMP | No transformat | -89 | 20.63169 | 6.10057 | 0.01008 | 0.05 | 1 | -163.39 | -14.6097 |
| AMP | MTC5 | NoAMP | Control | -24 | 20.63169 | 1.6451 | 0.98722 | 0.05 | 0 | -98.3903 | 50.39027 |
| AMP | MTC5 | NoAMP | Ligation insert | 15 | 20.63169 | 1.02819 | 0.99978 | 0.05 | 0 | -59.3903 | 89.39027 |
| AMP | MTC5 | NoAMP | CH3 | -4.33333 | 20.63169 | 0.29703 | 1 | 0.05 | 0 | -78.7236 | 70.05694 |
| AMP | MTC5 | NoAMP | MTC5 | -24 | 20.63169 | 1.6451 | 0.98722 | 0.05 | 0 | -98.3903 | 50.39027 |
| AMP | MTC5 | NoAMP | AdoHcy | 6.66667 | 20.63169 | 0.45697 | 1 | 0.05 | 0 | -67.7236 | 81.05694 |
| AMP | MTC5 | AMP | No transformat | 183 | 20.63169 | 12.54386 | 2.30E-07 | 0.05 | 1 | 108.6097 | 257.3903 |
| AMP | MTC5 | AMP | Control | -39.6667 | 20.63169 | 2.71898 | 0.7354 | 0.05 | 0 | -114.057 | 34.72361 |
| AMP | MTC5 | AMP | Ligation insert | -31 | 20.63169 | 2.12492 | 0.92574 | 0.05 | 0 | -105.39 | 43.39027 |
| AMP | MTC5 | AMP | CH3 | -22 | 20.63169 | 1.50801 | 0.99355 | 0.05 | 0 | -96.3903 | 52.39027 |
| AMP | AdoHcy | NoAMP | No transformat | -272 | 20.63169 | 18.64443 | 2.66E-07 | 0.05 | 1 | -346.39 | -197.61 |
| AMP | AdoHcy | NoAMP | Control | -207 | 20.63169 | 14.18896 | 5.15E-08 | 0.05 | 1 | -281.39 | -132.61 |
| AMP | AdoHcy | NoAMP | Ligation insert | -168 | 20.63169 | 11.51568 | 1.23E-06 | 0.05 | 1 | -242.39 | -93.6097 |
| AMP | AdoHcy | NoAMP | CH3 | -187.333 | 20.63169 | 12.84089 | 1.34E-07 | 0.05 | 1 | -261.724 | -112.943 |
| AMP | AdoHcy | NoAMP | MTC5 | -207 | 20.63169 | 14.18896 | 5.15E-08 | 0.05 | 1 | -281.39 | -132.61 |
| AMP | AdoHcy | NoAMP | AdoHcy | -176.333 | 20.63169 | 12.08689 | 4.90E-07 | 0.05 | 1 | -250.724 | -101.943 |
| AMP | AdoHcy | AMP | No transformat | 0 | 20.63169 | 0 | 1 | 0.05 | 0 | -74.3903 | 74.39027 |
| AMP | AdoHcy | AMP | Control | -222.667 | 20.63169 | 15.26284 | 3.18E-08 | 0.05 | 1 | -297.057 | -148.276 |
| AMP | AdoHcy | AMP | Ligation insert | -214 | 20.63169 | 14.66878 | 3.96E-08 | 0.05 | 1 | -288.39 | -139.61 |
| AMP | AdoHcy | AMP | CH3 | -205 | 20.63169 | 14.05187 | 5.69E-08 | 0.05 | 1 | -279.39 | -130.61 |
| AMP | AdoHcy | AMP | MTC5 | -183 | 20.63169 | 12.54386 | 2.30E-07 | 0.05 | 1 | -257.39 | -108.61 |

**Table S7. Two-way ANOVA results of DH5α with and without modified plasmids on control and kanamycin plates (statistically significant results highlighted in red).**

| Antibiotic | Modificatio | Antibiotic | Modificatio | MeanDiff | SEM | q Value | Prob | Alpha | Sig | LCL | UCL |
|---|---|---|---|---|---|---|---|---|---|---|---|
| NoKan | Control | NoKan | No transfor | -31.5 | 21.94216 | 2.03023 | 0.93336 | 0.05 | 0 | -118.614 | 55.61375 |
| NoKan | Ligation ins | NoKan | No transfor | -40 | 21.94216 | 2.57808 | 0.78116 | 0.05 | 0 | -127.114 | 47.11375 |
| NoKan | Ligation ins | NoKan | Control | -8.5 | 21.94216 | 0.54784 | 1 | 0.05 | 0 | -95.6138 | 78.61375 |
| NoKan | CH3 | NoKan | No transfor | -75 | 21.94216 | 4.83389 | 0.1178 | 0.05 | 0 | -162.114 | 12.11375 |
| NoKan | CH3 | NoKan | Control | -43.5 | 21.94216 | 2.80366 | 0.69721 | 0.05 | 0 | -130.614 | 43.61375 |
| NoKan | CH3 | NoKan | Ligation ins | -35 | 21.94216 | 2.25582 | 0.8819 | 0.05 | 0 | -122.114 | 52.11375 |
| NoKan | MTC5 | NoKan | No transfor | -33 | 21.94216 | 2.12691 | 0.91342 | 0.05 | 0 | -120.114 | 54.11375 |
| NoKan | MTC5 | NoKan | Control | -1.5 | 21.94216 | 0.09668 | 1 | 0.05 | 0 | -88.6138 | 85.61375 |
| NoKan | MTC5 | NoKan | Ligation ins | 7 | 21.94216 | 0.45116 | 1 | 0.05 | 0 | -80.1138 | 94.11375 |
| NoKan | MTC5 | NoKan | CH3 | 42 | 21.94216 | 2.70698 | 0.73411 | 0.05 | 0 | -45.1138 | 129.1138 |
| NoKan | AdoHcy | NoKan | No transfor | -79.5 | 21.94216 | 5.12393 | 0.08603 | 0.05 | 0 | -166.614 | 7.61375 |
| NoKan | AdoHcy | NoKan | Control | -48 | 21.94216 | 3.09369 | 0.58292 | 0.05 | 0 | -135.114 | 39.11375 |
| NoKan | AdoHcy | NoKan | Ligation ins | -39.5 | 21.94216 | 2.54585 | 0.79243 | 0.05 | 0 | -126.614 | 47.61375 |
| NoKan | AdoHcy | NoKan | CH3 | -4.5 | 21.94216 | 0.29003 | 1 | 0.05 | 0 | -91.6138 | 82.61375 |
| NoKan | AdoHcy | NoKan | MTC5 | -46.5 | 21.94216 | 2.99701 | 0.62118 | 0.05 | 0 | -133.614 | 40.61375 |
| Kan | No transfor | NoKan | No transfor | -297.5 | 21.94216 | 19.17444 | 4.20E-07 | 0.05 | 1 | -384.614 | -210.386 |
| Kan | No transfor | NoKan | Control | -266 | 21.94216 | 17.1442 | 1.97E-06 | 0.05 | 1 | -353.114 | -178.886 |
| Kan | No transfor | NoKan | Ligation ins | -257.5 | 21.94216 | 16.59636 | 4.11E-06 | 0.05 | 1 | -344.614 | -170.386 |
| Kan | No transfor | NoKan | CH3 | -222.5 | 21.94216 | 14.34055 | 1.25E-05 | 0.05 | 1 | -309.614 | -135.386 |
| Kan | No transfor | NoKan | MTC5 | -264.5 | 21.94216 | 17.04752 | 3.54E-06 | 0.05 | 1 | -351.614 | -177.386 |
| Kan | No transfor | NoKan | AdoHcy | -218 | 21.94216 | 14.05051 | 1.54E-05 | 0.05 | 1 | -305.114 | -130.886 |
| Kan | Control | NoKan | No transfor | -64.5 | 21.94216 | 4.15715 | 0.23672 | 0.05 | 0 | -151.614 | 22.61375 |
| Kan | Control | NoKan | Control | -33 | 21.94216 | 2.12691 | 0.91342 | 0.05 | 0 | -120.114 | 54.11375 |
| Kan | Control | NoKan | Ligation ins | -24.5 | 21.94216 | 1.57907 | 0.98718 | 0.05 | 0 | -111.614 | 62.61375 |
| Kan | Control | NoKan | CH3 | 10.5 | 21.94216 | 0.67674 | 0.99999 | 0.05 | 0 | -76.6138 | 97.61375 |
| Kan | Control | NoKan | MTC5 | -31.5 | 21.94216 | 2.03023 | 0.93336 | 0.05 | 0 | -118.614 | 55.61375 |
| Kan | Control | NoKan | AdoHcy | 15 | 21.94216 | 0.96678 | 0.99979 | 0.05 | 0 | -72.1138 | 102.1138 |
| Kan | Control | Kan | No transfor | 233 | 21.94216 | 15.01729 | 7.74E-06 | 0.05 | 1 | 145.8863 | 320.1138 |
| Kan | Ligation ins | NoKan | No transfor | -28 | 21.94216 | 1.80465 | 0.96772 | 0.05 | 0 | -115.114 | 59.11375 |
| Kan | Ligation ins | NoKan | Control | 3.5 | 21.94216 | 0.22558 | 1 | 0.05 | 0 | -83.6138 | 90.61375 |
| Kan | Ligation ins | NoKan | Ligation ins | 12 | 21.94216 | 0.77342 | 0.99998 | 0.05 | 0 | -75.1138 | 99.11375 |
| Kan | Ligation ins | NoKan | CH3 | 47 | 21.94216 | 3.02924 | 0.60841 | 0.05 | 0 | -40.1138 | 134.1138 |
| Kan | Ligation ins | NoKan | MTC5 | 5 | 21.94216 | 0.32226 | 1 | 0.05 | 0 | -82.1138 | 92.11375 |
| Kan | Ligation ins | NoKan | AdoHcy | 51.5 | 21.94216 | 3.31927 | 0.49558 | 0.05 | 0 | -35.6138 | 138.6138 |
| Kan | Ligation ins | Kan | No transfor | 269.5 | 21.94216 | 17.36978 | 1.73E-06 | 0.05 | 1 | 182.3863 | 356.6138 |
| Kan | Ligation ins | Kan | Control | 36.5 | 21.94216 | 2.35249 | 0.85474 | 0.05 | 0 | -50.6138 | 123.6138 |
| Kan | CH3 | NoKan | No transfor | -56 | 21.94216 | 3.60931 | 0.39206 | 0.05 | 0 | -143.114 | 31.11375 |
| Kan | CH3 | NoKan | Control | -24.5 | 21.94216 | 1.57907 | 0.98718 | 0.05 | 0 | -111.614 | 62.61375 |
| Kan | CH3 | NoKan | Ligation ins | -16 | 21.94216 | 1.03123 | 0.99962 | 0.05 | 0 | -103.114 | 71.11375 |
| Kan | CH3 | NoKan | CH3 | 19 | 21.94216 | 1.22459 | 0.99829 | 0.05 | 0 | -68.1138 | 106.1138 |
| Kan | CH3 | NoKan | MTC5 | -23 | 21.94216 | 1.48239 | 0.99199 | 0.05 | 0 | -110.114 | 64.11375 |
| Kan | CH3 | NoKan | AdoHcy | 23.5 | 21.94216 | 1.51462 | 0.99058 | 0.05 | 0 | -63.6138 | 110.6138 |
| Kan | CH3 | Kan | No transfor | 241.5 | 21.94216 | 15.56513 | 5.30E-06 | 0.05 | 1 | 154.3863 | 328.6138 |
| Kan | CH3 | Kan | Control | 8.5 | 21.94216 | 0.54784 | 1 | 0.05 | 0 | -78.6138 | 95.61375 |
| Kan | CH3 | Kan | Ligation ins | -28 | 21.94216 | 1.80465 | 0.96772 | 0.05 | 0 | -115.114 | 59.11375 |
| Kan | MTC5 | NoKan | No transfor | -85 | 21.94216 | 5.47841 | 0.05818 | 0.05 | 0 | -172.114 | 2.11375 |
| Kan | MTC5 | NoKan | Control | -53.5 | 21.94216 | 3.44818 | 0.44804 | 0.05 | 0 | -140.614 | 33.61375 |
| Kan | MTC5 | NoKan | Ligation ins | -45 | 21.94216 | 2.90033 | 0.65942 | 0.05 | 0 | -132.114 | 42.11375 |
| Kan | MTC5 | NoKan | CH3 | -10 | 21.94216 | 0.64452 | 1 | 0.05 | 0 | -97.1138 | 77.11375 |
| Kan | MTC5 | NoKan | MTC5 | -52 | 21.94216 | 3.3515 | 0.48349 | 0.05 | 0 | -139.114 | 35.11375 |
| Kan | MTC5 | NoKan | AdoHcy | -5.5 | 21.94216 | 0.35449 | 1 | 0.05 | 0 | -92.6138 | 81.61375 |
| Kan | MTC5 | Kan | No transfor | 212.5 | 21.94216 | 13.69603 | 2.02E-05 | 0.05 | 1 | 125.3863 | 299.6138 |
| Kan | MTC5 | Kan | Control | -20.5 | 21.94216 | 1.32126 | 0.99678 | 0.05 | 0 | -107.614 | 66.61375 |
| Kan | MTC5 | Kan | Ligation ins | -57 | 21.94216 | 3.67376 | 0.37089 | 0.05 | 0 | -144.114 | 30.11375 |
| Kan | MTC5 | Kan | CH3 | -29 | 21.94216 | 1.8691 | 0.95957 | 0.05 | 0 | -116.114 | 58.11375 |
| Kan | AdoHcy | NoKan | No transfor | -297.5 | 21.94216 | 19.17444 | 4.20E-07 | 0.05 | 1 | -384.614 | -210.386 |
| Kan | AdoHcy | NoKan | Control | -266 | 21.94216 | 17.1442 | 1.97E-06 | 0.05 | 1 | -353.114 | -178.886 |
| Kan | AdoHcy | NoKan | Ligation ins | -257.5 | 21.94216 | 16.59636 | 4.11E-06 | 0.05 | 1 | -344.614 | -170.386 |
| Kan | AdoHcy | NoKan | CH3 | -222.5 | 21.94216 | 14.34055 | 1.25E-05 | 0.05 | 1 | -309.614 | -135.386 |
| Kan | AdoHcy | NoKan | MTC5 | -264.5 | 21.94216 | 17.04752 | 3.54E-06 | 0.05 | 1 | -351.614 | -177.386 |
| Kan | AdoHcy | NoKan | AdoHcy | -218 | 21.94216 | 14.05051 | 1.54E-05 | 0.05 | 1 | -305.114 | -130.886 |
| Kan | AdoHcy | Kan | No transfor | 0 | 21.94216 | 0 | 1 | 0.05 | 0 | -87.1138 | 87.11375 |
| Kan | AdoHcy | Kan | Control | -233 | 21.94216 | 15.01729 | 7.74E-06 | 0.05 | 1 | -320.114 | -145.886 |
| Kan | AdoHcy | Kan | Ligation ins | -269.5 | 21.94216 | 17.36978 | 1.73E-06 | 0.05 | 1 | -356.614 | -182.386 |
| Kan | AdoHcy | Kan | CH3 | -241.5 | 21.94216 | 15.56513 | 5.30E-06 | 0.05 | 1 | -328.614 | -154.386 |
| Kan | AdoHcy | Kan | MTC5 | -212.5 | 21.94216 | 13.69603 | 2.02E-05 | 0.05 | 1 | -299.614 | -125.386 |

**Table S8. Two-way ANOVA results of BL21 with and without modified plasmids on control and ampicillin plates (statistically significant results highlighted in red).**

| Antibiotic | Modificatio | Antibiotic | Modificatio | MeanDiff | SEM | q Value | Prob | Alpha | Sig | LCL | UCL |
|---|---|---|---|---|---|---|---|---|---|---|---|
| NoAMP | Control | NoAMP | No transfor | -87.5 | 29.68024 | 4.16923 | 0.23393 | 0.05 | 0 | -205.335 | 30.33512 |
| NoAMP | Ligation ins | NoAMP | No transfor | -83 | 29.68024 | 3.95481 | 0.2875 | 0.05 | 0 | -200.835 | 34.83512 |
| NoAMP | Ligation ins | NoAMP | Control | 4.5 | 29.68024 | 0.21442 | 1 | 0.05 | 0 | -113.335 | 122.3351 |
| NoAMP | CH3 | NoAMP | No transfor | -101.5 | 29.68024 | 4.8363 | 0.11749 | 0.05 | 0 | -219.335 | 16.33512 |
| NoAMP | CH3 | NoAMP | Control | -14 | 29.68024 | 0.66708 | 0.99999 | 0.05 | 0 | -131.835 | 103.8351 |
| NoAMP | CH3 | NoAMP | Ligation ins | -18.5 | 29.68024 | 0.88149 | 0.99991 | 0.05 | 0 | -136.335 | 99.33512 |
| NoAMP | MTC5 | NoAMP | No transfor | -107 | 29.68024 | 5.09837 | 0.08847 | 0.05 | 0 | -224.835 | 10.83512 |
| NoAMP | MTC5 | NoAMP | Control | -19.5 | 29.68024 | 0.92914 | 0.99986 | 0.05 | 0 | -137.335 | 98.33512 |
| NoAMP | MTC5 | NoAMP | Ligation ins | -24 | 29.68024 | 1.14356 | 0.99905 | 0.05 | 0 | -141.835 | 93.83512 |
| NoAMP | MTC5 | NoAMP | CH3 | -5.5 | 29.68024 | 0.26207 | 1 | 0.05 | 0 | -123.335 | 112.3351 |
| NoAMP | AdoHcy | NoAMP | No transfor | -92 | 29.68024 | 4.38365 | 0.18875 | 0.05 | 0 | -209.835 | 25.83512 |
| NoAMP | AdoHcy | NoAMP | Control | -4.5 | 29.68024 | 0.21442 | 1 | 0.05 | 0 | -122.335 | 113.3351 |
| NoAMP | AdoHcy | NoAMP | Ligation ins | -9 | 29.68024 | 0.42883 | 1 | 0.05 | 0 | -126.835 | 108.8351 |
| NoAMP | AdoHcy | NoAMP | CH3 | 9.5 | 29.68024 | 0.45266 | 1 | 0.05 | 0 | -108.335 | 127.3351 |
| NoAMP | AdoHcy | NoAMP | MTC5 | 15 | 29.68024 | 0.71472 | 0.99999 | 0.05 | 0 | -102.835 | 132.8351 |
| AMP | No transfor | NoAMP | No transfor | -334.5 | 29.68024 | 15.93836 | 4.14E-06 | 0.05 | 1 | -452.335 | -216.665 |
| AMP | No transfor | NoAMP | Control | -247 | 29.68024 | 11.76913 | 9.73E-05 | 0.05 | 1 | -364.835 | -129.165 |
| AMP | No transfor | NoAMP | Ligation ins | -251.5 | 29.68024 | 11.98355 | 8.11E-05 | 0.05 | 1 | -369.335 | -133.665 |
| AMP | No transfor | NoAMP | CH3 | -233 | 29.68024 | 11.10206 | 1.75E-04 | 0.05 | 1 | -350.835 | -115.165 |
| AMP | No transfor | NoAMP | MTC5 | -227.5 | 29.68024 | 10.83999 | 2.21E-04 | 0.05 | 1 | -345.335 | -109.665 |
| AMP | No transfor | NoAMP | AdoHcy | -242.5 | 29.68024 | 11.55472 | 1.17E-04 | 0.05 | 1 | -360.335 | -124.665 |
| AMP | Control | NoAMP | No transfor | -84.5 | 29.68024 | 4.02628 | 0.26868 | 0.05 | 0 | -202.335 | 33.33512 |
| AMP | Control | NoAMP | Control | 3 | 29.68024 | 0.14294 | 1 | 0.05 | 0 | -114.835 | 120.8351 |
| AMP | Control | NoAMP | Ligation ins | -1.5 | 29.68024 | 0.07147 | 1 | 0.05 | 0 | -119.335 | 116.3351 |
| AMP | Control | NoAMP | CH3 | 17 | 29.68024 | 0.81002 | 0.99996 | 0.05 | 0 | -100.835 | 134.8351 |
| AMP | Control | NoAMP | MTC5 | 22.5 | 29.68024 | 1.07209 | 0.99947 | 0.05 | 0 | -95.3351 | 140.3351 |
| AMP | Control | NoAMP | AdoHcy | 7.5 | 29.68024 | 0.35736 | 1 | 0.05 | 0 | -110.335 | 125.3351 |
| AMP | Control | AMP | No transfor | 250 | 29.68024 | 11.91208 | 8.61E-05 | 0.05 | 1 | 132.1649 | 367.8351 |
| AMP | Ligation ins | NoAMP | No transfor | -80.5 | 29.68024 | 3.83569 | 0.32102 | 0.05 | 0 | -198.335 | 37.33512 |
| AMP | Ligation ins | NoAMP | Control | 7 | 29.68024 | 0.33354 | 1 | 0.05 | 0 | -110.835 | 124.8351 |
| AMP | Ligation ins | NoAMP | Ligation ins | 2.5 | 29.68024 | 0.11912 | 1 | 0.05 | 0 | -115.335 | 120.3351 |
| AMP | Ligation ins | NoAMP | CH3 | 21 | 29.68024 | 1.00061 | 0.99971 | 0.05 | 0 | -96.8351 | 138.8351 |
| AMP | Ligation ins | NoAMP | MTC5 | 26.5 | 29.68024 | 1.26268 | 0.99778 | 0.05 | 0 | -91.3351 | 144.3351 |
| AMP | Ligation ins | NoAMP | AdoHcy | 11.5 | 29.68024 | 0.54796 | 1 | 0.05 | 0 | -106.335 | 129.3351 |
| AMP | Ligation ins | AMP | No transfor | 254 | 29.68024 | 12.10267 | 7.34E-05 | 0.05 | 1 | 136.1649 | 371.8351 |
| AMP | Ligation ins | AMP | Control | 4 | 29.68024 | 0.19059 | 1 | 0.05 | 0 | -113.835 | 121.8351 |
| AMP | CH3 | NoAMP | No transfor | -76 | 29.68024 | 3.62127 | 0.38807 | 0.05 | 0 | -193.835 | 41.83512 |
| AMP | CH3 | NoAMP | Control | 11.5 | 29.68024 | 0.54796 | 1 | 0.05 | 0 | -106.335 | 129.3351 |
| AMP | CH3 | NoAMP | Ligation ins | 7 | 29.68024 | 0.33354 | 1 | 0.05 | 0 | -110.835 | 124.8351 |
| AMP | CH3 | NoAMP | CH3 | 25.5 | 29.68024 | 1.21503 | 0.9984 | 0.05 | 0 | -92.3351 | 143.3351 |
| AMP | CH3 | NoAMP | MTC5 | 31 | 29.68024 | 1.4771 | 0.99221 | 0.05 | 0 | -86.8351 | 148.8351 |
| AMP | CH3 | NoAMP | AdoHcy | 16 | 29.68024 | 0.76237 | 0.99998 | 0.05 | 0 | -101.835 | 133.8351 |
| AMP | CH3 | AMP | No transfor | 258.5 | 29.68024 | 12.31709 | 6.12E-05 | 0.05 | 1 | 140.6649 | 376.3351 |
| AMP | CH3 | AMP | Control | 8.5 | 29.68024 | 0.40501 | 1 | 0.05 | 0 | -109.335 | 126.3351 |
| AMP | CH3 | AMP | Ligation ins | 4.5 | 29.68024 | 0.21442 | 1 | 0.05 | 0 | -113.335 | 122.3351 |
| AMP | MTC5 | NoAMP | No transfor | -100 | 29.68024 | 4.76483 | 0.12682 | 0.05 | 0 | -217.835 | 17.83512 |
| AMP | MTC5 | NoAMP | Control | -12.5 | 29.68024 | 0.5956 | 1 | 0.05 | 0 | -130.335 | 105.3351 |
| AMP | MTC5 | NoAMP | Ligation ins | -17 | 29.68024 | 0.81002 | 0.99996 | 0.05 | 0 | -134.835 | 100.8351 |
| AMP | MTC5 | NoAMP | CH3 | 1.5 | 29.68024 | 0.07147 | 1 | 0.05 | 0 | -116.335 | 119.3351 |
| AMP | MTC5 | NoAMP | MTC5 | 7 | 29.68024 | 0.33354 | 1 | 0.05 | 0 | -110.835 | 124.8351 |
| AMP | MTC5 | NoAMP | AdoHcy | -8 | 29.68024 | 0.38119 | 1 | 0.05 | 0 | -125.835 | 109.8351 |
| AMP | MTC5 | AMP | No transfor | 234.5 | 29.68024 | 11.17353 | 1.64E-04 | 0.05 | 1 | 116.6649 | 352.3351 |
| AMP | MTC5 | AMP | Control | -15.5 | 29.68024 | 0.73855 | 0.99998 | 0.05 | 0 | -133.335 | 102.3351 |
| AMP | MTC5 | AMP | Ligation ins | -19.5 | 29.68024 | 0.92914 | 0.99986 | 0.05 | 0 | -137.335 | 98.33512 |
| AMP | MTC5 | AMP | CH3 | -24 | 29.68024 | 1.14356 | 0.99905 | 0.05 | 0 | -141.835 | 93.83512 |
| AMP | AdoHcy | NoAMP | No transfor | -334.5 | 29.68024 | 15.93836 | 4.14E-06 | 0.05 | 1 | -452.335 | -216.665 |
| AMP | AdoHcy | NoAMP | Control | -247 | 29.68024 | 11.76913 | 9.73E-05 | 0.05 | 1 | -364.835 | -129.165 |
| AMP | AdoHcy | NoAMP | Ligation ins | -251.5 | 29.68024 | 11.98355 | 8.11E-05 | 0.05 | 1 | -369.335 | -133.665 |
| AMP | AdoHcy | NoAMP | CH3 | -233 | 29.68024 | 11.10206 | 1.75E-04 | 0.05 | 1 | -350.835 | -115.165 |
| AMP | AdoHcy | NoAMP | MTC5 | -227.5 | 29.68024 | 10.83999 | 2.21E-04 | 0.05 | 1 | -345.335 | -109.665 |
| AMP | AdoHcy | NoAMP | AdoHcy | -242.5 | 29.68024 | 11.55472 | 1.17E-04 | 0.05 | 1 | -360.335 | -124.665 |
| AMP | AdoHcy | AMP | No transfor | 0 | 29.68024 | 0 | 1 | 0.05 | 0 | -117.835 | 117.8351 |
| AMP | AdoHcy | AMP | Control | -250 | 29.68024 | 11.91208 | 8.61E-05 | 0.05 | 1 | -367.835 | -132.165 |
| AMP | AdoHcy | AMP | Ligation ins | -254 | 29.68024 | 12.10267 | 7.34E-05 | 0.05 | 1 | -371.835 | -136.165 |
| AMP | AdoHcy | AMP | CH3 | -258.5 | 29.68024 | 12.31709 | 6.12E-05 | 0.05 | 1 | -376.335 | -140.665 |
| AMP | AdoHcy | AMP | MTC5 | -234.5 | 29.68024 | 11.17353 | 1.64E-04 | 0.05 | 1 | -352.335 | -116.665 |

**Table S9. Two-way ANOVA results of BL21 with and without modified plasmids on control and kanamycin plates (statistically significant results highlighted in red).**

| Antibiotic | Modificatio | Antibiotic | Modificatio | MeanDiff | SEM | q Value | Prob | Alpha | Sig | LCL | UCL |
|---|---|---|---|---|---|---|---|---|---|---|---|
| NoKan | Control | NoKan | No transfor | -58 | 23.7364 | 3.45564 | 0.44536 | 0.05 | 0 | -152.237 | 36.23716 |
| NoKan | Ligation ins | NoKan | No transfor | -73 | 23.7364 | 4.34934 | 0.19544 | 0.05 | 0 | -167.237 | 21.23716 |
| NoKan | Ligation ins | NoKan | Control | -15 | 23.7364 | 0.8937 | 0.9999 | 0.05 | 0 | -109.237 | 79.23716 |
| NoKan | CH3 | NoKan | No transfor | -96 | 23.7364 | 5.71968 | 0.04447 | 0.05 | 1 | -190.237 | -1.76284 |
| NoKan | CH3 | NoKan | Control | -38 | 23.7364 | 2.26404 | 0.87971 | 0.05 | 0 | -132.237 | 56.23716 |
| NoKan | CH3 | NoKan | Ligation ins | -23 | 23.7364 | 1.37034 | 0.99567 | 0.05 | 0 | -117.237 | 71.23716 |
| NoKan | MTC5 | NoKan | No transfor | -70.5 | 23.7364 | 4.20039 | 0.22686 | 0.05 | 0 | -164.737 | 23.73716 |
| NoKan | MTC5 | NoKan | Control | -12.5 | 23.7364 | 0.74475 | 0.99998 | 0.05 | 0 | -106.737 | 81.73716 |
| NoKan | MTC5 | NoKan | Ligation ins | 2.5 | 23.7364 | 0.14895 | 1 | 0.05 | 0 | -91.7372 | 96.73716 |
| NoKan | MTC5 | NoKan | CH3 | 25.5 | 23.7364 | 1.51929 | 0.99036 | 0.05 | 0 | -68.7372 | 119.7372 |
| NoKan | AdoHcy | NoKan | No transfor | -73 | 23.7364 | 4.34934 | 0.19544 | 0.05 | 0 | -167.237 | 21.23716 |
| NoKan | AdoHcy | NoKan | Control | -15 | 23.7364 | 0.8937 | 0.9999 | 0.05 | 0 | -109.237 | 79.23716 |
| NoKan | AdoHcy | NoKan | Ligation ins | 0 | 23.7364 | 0 | 1 | 0.05 | 0 | -94.2372 | 94.23716 |
| NoKan | AdoHcy | NoKan | CH3 | 23 | 23.7364 | 1.37034 | 0.99567 | 0.05 | 0 | -71.2372 | 117.2372 |
| NoKan | AdoHcy | NoKan | MTC5 | -2.5 | 23.7364 | 0.14895 | 1 | 0.05 | 0 | -96.7372 | 91.73716 |
| Kan | No transfor | NoKan | No transfor | -330.5 | 23.7364 | 19.69117 | 2.83E-07 | 0.05 | 1 | -424.737 | -236.263 |
| Kan | No transfor | NoKan | Control | -272.5 | 23.7364 | 16.23554 | 3.42E-06 | 0.05 | 1 | -366.737 | -178.263 |
| Kan | No transfor | NoKan | Ligation ins | -257.5 | 23.7364 | 15.34184 | 6.18E-06 | 0.05 | 1 | -351.737 | -163.263 |
| Kan | No transfor | NoKan | CH3 | -234.5 | 23.7364 | 13.9715 | 1.63E-05 | 0.05 | 1 | -328.737 | -140.263 |
| Kan | No transfor | NoKan | MTC5 | -260 | 23.7364 | 15.49079 | 5.59E-06 | 0.05 | 1 | -354.237 | -165.763 |
| Kan | No transfor | NoKan | AdoHcy | -257.5 | 23.7364 | 15.34184 | 6.18E-06 | 0.05 | 1 | -351.737 | -163.263 |
| Kan | Control | NoKan | No transfor | -79.5 | 23.7364 | 4.73661 | 0.13068 | 0.05 | 0 | -173.737 | 14.73716 |
| Kan | Control | NoKan | Control | -21.5 | 23.7364 | 1.28097 | 0.9975 | 0.05 | 0 | -115.737 | 72.73716 |
| Kan | Control | NoKan | Ligation ins | -6.5 | 23.7364 | 0.38727 | 1 | 0.05 | 0 | -100.737 | 87.73716 |
| Kan | Control | NoKan | CH3 | 16.5 | 23.7364 | 0.98307 | 0.99976 | 0.05 | 0 | -77.7372 | 110.7372 |
| Kan | Control | NoKan | MTC5 | -9 | 23.7364 | 0.53622 | 1 | 0.05 | 0 | -103.237 | 85.23716 |
| Kan | Control | NoKan | AdoHcy | -6.5 | 23.7364 | 0.38727 | 1 | 0.05 | 0 | -100.737 | 87.73716 |
| Kan | Control | Kan | No transfor | 251 | 23.7364 | 14.95457 | 8.09E-06 | 0.05 | 1 | 156.7628 | 345.2372 |
| Kan | Ligation ins | NoKan | No transfor | -82 | 23.7364 | 4.88556 | 0.11144 | 0.05 | 0 | -176.237 | 12.23716 |
| Kan | Ligation ins | NoKan | Control | -24 | 23.7364 | 1.42992 | 0.99394 | 0.05 | 0 | -118.237 | 70.23716 |
| Kan | Ligation ins | NoKan | Ligation ins | -9 | 23.7364 | 0.53622 | 1 | 0.05 | 0 | -103.237 | 85.23716 |
| Kan | Ligation ins | NoKan | CH3 | 14 | 23.7364 | 0.83412 | 0.99995 | 0.05 | 0 | -80.2372 | 108.2372 |
| Kan | Ligation ins | NoKan | MTC5 | -11.5 | 23.7364 | 0.68517 | 0.99999 | 0.05 | 0 | -105.737 | 82.73716 |
| Kan | Ligation ins | NoKan | AdoHcy | -9 | 23.7364 | 0.53622 | 1 | 0.05 | 0 | -103.237 | 85.23716 |
| Kan | Ligation ins | Kan | No transfor | 248.5 | 23.7364 | 14.80562 | 8.99E-06 | 0.05 | 1 | 154.2628 | 342.7372 |
| Kan | Ligation ins | Kan | Control | -2.5 | 23.7364 | 0.14895 | 1 | 0.05 | 0 | -96.7372 | 91.73716 |
| Kan | CH3 | NoKan | No transfor | -102.5 | 23.7364 | 6.10695 | 0.02884 | 0.05 | 1 | -196.737 | -8.26284 |
| Kan | CH3 | NoKan | Control | -44.5 | 23.7364 | 2.65131 | 0.75478 | 0.05 | 0 | -138.737 | 49.73716 |
| Kan | CH3 | NoKan | Ligation ins | -29.5 | 23.7364 | 1.75761 | 0.97289 | 0.05 | 0 | -123.737 | 64.73716 |
| Kan | CH3 | NoKan | CH3 | -6.5 | 23.7364 | 0.38727 | 1 | 0.05 | 0 | -100.737 | 87.73716 |
| Kan | CH3 | NoKan | MTC5 | -32 | 23.7364 | 1.90656 | 0.95424 | 0.05 | 0 | -126.237 | 62.23716 |
| Kan | CH3 | NoKan | AdoHcy | -29.5 | 23.7364 | 1.75761 | 0.97289 | 0.05 | 0 | -123.737 | 64.73716 |
| Kan | CH3 | Kan | No transfor | 228 | 23.7364 | 13.58423 | 2.20E-05 | 0.05 | 1 | 133.7628 | 322.2372 |
| Kan | CH3 | Kan | Control | -23 | 23.7364 | 1.37034 | 0.99567 | 0.05 | 0 | -117.237 | 71.23716 |
| Kan | CH3 | Kan | Ligation ins | -20.5 | 23.7364 | 1.22139 | 0.99833 | 0.05 | 0 | -114.737 | 73.73716 |
| Kan | MTC5 | NoKan | No transfor | -113.5 | 23.7364 | 6.76232 | 0.01388 | 0.05 | 1 | -207.737 | -19.2628 |
| Kan | MTC5 | NoKan | Control | -55.5 | 23.7364 | 3.30669 | 0.50033 | 0.05 | 0 | -149.737 | 38.73716 |
| Kan | MTC5 | NoKan | Ligation ins | -40.5 | 23.7364 | 2.41299 | 0.83631 | 0.05 | 0 | -134.737 | 53.73716 |
| Kan | MTC5 | NoKan | CH3 | -17.5 | 23.7364 | 1.04265 | 0.99958 | 0.05 | 0 | -111.737 | 76.73716 |
| Kan | MTC5 | NoKan | MTC5 | -43 | 23.7364 | 2.56194 | 0.78683 | 0.05 | 0 | -137.237 | 51.23716 |
| Kan | MTC5 | NoKan | AdoHcy | -40.5 | 23.7364 | 2.41299 | 0.83631 | 0.05 | 0 | -134.737 | 53.73716 |
| Kan | MTC5 | Kan | No transfor | 217 | 23.7364 | 12.92885 | 3.70E-05 | 0.05 | 1 | 122.7628 | 311.2372 |
| Kan | MTC5 | Kan | Control | -34 | 23.7364 | 2.02572 | 0.93421 | 0.05 | 0 | -128.237 | 60.23716 |
| Kan | MTC5 | Kan | Ligation ins | -31.5 | 23.7364 | 1.87677 | 0.95852 | 0.05 | 0 | -125.737 | 62.73716 |
| Kan | MTC5 | Kan | CH3 | -11 | 23.7364 | 0.65538 | 0.99999 | 0.05 | 0 | -105.237 | 83.23716 |
| Kan | AdoHcy | NoKan | No transfor | -330.5 | 23.7364 | 19.69117 | 2.83E-07 | 0.05 | 1 | -424.737 | -236.263 |
| Kan | AdoHcy | NoKan | Control | -272.5 | 23.7364 | 16.23554 | 3.42E-06 | 0.05 | 1 | -366.737 | -178.263 |
| Kan | AdoHcy | NoKan | Ligation ins | -257.5 | 23.7364 | 15.34184 | 6.18E-06 | 0.05 | 1 | -351.737 | -163.263 |
| Kan | AdoHcy | NoKan | CH3 | -234.5 | 23.7364 | 13.9715 | 1.63E-05 | 0.05 | 1 | -328.737 | -140.263 |
| Kan | AdoHcy | NoKan | MTC5 | -260 | 23.7364 | 15.49079 | 5.59E-06 | 0.05 | 1 | -354.237 | -165.763 |
| Kan | AdoHcy | NoKan | AdoHcy | -257.5 | 23.7364 | 15.34184 | 6.18E-06 | 0.05 | 1 | -351.737 | -163.263 |
| Kan | AdoHcy | Kan | No transfor | 0 | 23.7364 | 0 | 1 | 0.05 | 0 | -94.2372 | 94.23716 |
| Kan | AdoHcy | Kan | Control | -251 | 23.7364 | 14.95457 | 8.09E-06 | 0.05 | 1 | -345.237 | -156.763 |
| Kan | AdoHcy | Kan | Ligation ins | -248.5 | 23.7364 | 14.80562 | 8.99E-06 | 0.05 | 1 | -342.737 | -154.263 |
| Kan | AdoHcy | Kan | CH3 | -228 | 23.7364 | 13.58423 | 2.20E-05 | 0.05 | 1 | -322.237 | -133.763 |
| Kan | AdoHcy | Kan | MTC5 | -217 | 23.7364 | 12.92885 | 3.70E-05 | 0.05 | 1 | -311.237 | -122.763 |

**Table S10. Two-way ANOVA results of BL21 and DH5α transformed with and without modified eGFP-AmpR containing plasmids on ampicillin plates (statistically significant results highlighted in red).**

| Bacterial subspecies | Plasmid modification | Bacterial subspecies | Plasmid modification | MeanDiff | SEM | q Value | Prob | Alpha | Sig | LCL | UCL |
|---|---|---|---|---|---|---|---|---|---|---|---|
| DH5a | No transformation | DH5a | Control | -269 | 17.96602 | 21.17461 | 2.55E-07 | 0.05 | 1 | -329.346 | -208.654 |
| DH5a | AdoHcy-azide | DH5a | Control | -29.6667 | 17.96602 | 2.33524 | 0.58399 | 0.05 | 0 | -90.0131 | 30.67975 |
| DH5a | AdoHcy-azide | DH5a | No transformation | 239.3333 | 17.96602 | 18.83937 | 1.88E-07 | 0.05 | 1 | 178.9869 | 299.6798 |
| BL21 | Control | DH5a | Control | -0.66667 | 17.96602 | 0.05248 | 1 | 0.05 | 0 | -61.0131 | 59.67975 |
| BL21 | Control | DH5a | No transformation | 268.3333 | 17.96602 | 21.12213 | 2.55E-07 | 0.05 | 1 | 207.9869 | 328.6798 |
| BL21 | Control | DH5a | AdoHcy-azide | 29 | 17.96602 | 2.28276 | 0.6053 | 0.05 | 0 | -31.3464 | 89.34642 |
| BL21 | No transformation | DH5a | Control | -269 | 17.96602 | 21.17461 | 2.55E-07 | 0.05 | 1 | -329.346 | -208.654 |
| BL21 | No transformation | DH5a | No transformation | 0 | 17.96602 | 0 | 1 | 0.05 | 0 | -60.3464 | 60.34642 |
| BL21 | No transformation | DH5a | AdoHcy-azide | -239.333 | 17.96602 | 18.83937 | 1.88E-07 | 0.05 | 1 | -299.68 | -178.987 |
| BL21 | No transformation | BL21 | Control | -268.333 | 17.96602 | 21.12213 | 2.55E-07 | 0.05 | 1 | -328.68 | -207.987 |
| BL21 | AdoHcy-azide | DH5a | Control | -47.6667 | 17.96602 | 3.75213 | 0.15724 | 0.05 | 0 | -108.013 | 12.67975 |
| BL21 | AdoHcy-azide | DH5a | No transformation | 221.3333 | 17.96602 | 17.42248 | 4.33E-07 | 0.05 | 1 | 160.9869 | 281.6798 |
| BL21 | AdoHcy-azide | DH5a | AdoHcy-azide | -18 | 17.96602 | 1.41689 | 0.90863 | 0.05 | 0 | -78.3464 | 42.34642 |
| BL21 | AdoHcy-azide | BL21 | Control | -47 | 17.96602 | 3.69965 | 0.16649 | 0.05 | 0 | -107.346 | 13.34642 |
| BL21 | AdoHcy-azide | BL21 | No transformation | 221.3333 | 17.96602 | 17.42248 | 4.33E-07 | 0.05 | 1 | 160.9869 | 281.6798 |
